# Effect of set up protocols on the accuracy of alchemical free energy calculation over a set of ACK1 inhibitors

**DOI:** 10.1101/333120

**Authors:** José M. Granadino-Roldán, Antonia S. J. S. Mey, Juan J. Pérez González, Stefano Bosisio, Jaime Rubio-Martinez, Julien Michel

## Abstract

Hit-to-lead virtual screening frequently relies on a cascade of computational methods that starts with rapid calculations applied to a large number of compounds and ends with more expensive computations restricted to a subset of compounds that passed initial filters. This work focuses on set up protocols for alchemical free energy (AFE) scoring in the context of a Docking – MM/PBSA – AFE cascade. A dataset of 15 congeneric inhibitors of the ACK1 protein was used to evaluate the performance of AFE set up protocols that varied in the steps taken to prepare input files (using previously docked and best scored poses, manual selection of poses, manual placement of binding site water molecules). The main finding is that use of knowledge derived from X-ray structures to model binding modes, together with the manual placement of a bridging water molecule, improves the R^2^ from 0.45 ± 0.06 to 0.76 ± 0.02 and decreases the mean unsigned error from 2.11 ± 0.08 to 1.24 ± 0.04 kcal mol^-1^. By contrast a brute force automated protocol that increased the sampling time ten-fold lead to little improvements in accuracy. Besides, it is shown that for the present dataset hysteresis can be used to flag poses that need further attention even without prior knowledge of experimental binding affinities.

## INTRODUCTION

There is continuous interest in computational methods to decrease time and costs of hit-to-lead and lead optimization efforts in preclinical drug discovery [1]. A recurring topic in computational chemistry is the use of virtual *in silico* screens to find ligands for proteins [2, 3]. Typically, the goal is to filter via a cascade of computational methods a large library to focus experimental efforts on a small number of molecules. Usually inexpensive methodologies are applied first to eliminate a large number of poorly suited molecules, with more expensive calculations reserved to a subset of promising ligands. This approach may be applied in the context of hit discovery where the goal is to identify a few weak binders from a library of existing molecules; or for hit-to-lead efforts where the goal is to identify analogues of a hit structure that could be prioritized for synthesis and assays. In both cases the main steps frequently involve library screening, docking, initial scoring, and re-scoring with diverse molecular simulation methods such as Molecular Mechanics Poisson Boltzmann (Generalized Born) Surface Area (MM/PBSA) [4], Linear Interaction Energy (LIE) [5] or Free energy Perturbation (FEP) [6] methods [7].

In a previous study a multistep docking and scoring protocol was benchmarked in the context of re-scoring with the MM/PB(GB)SA method [8]. The set of ligands analysed in that study belonged to the same scaffold and it was assumed that the core binding mode of the conserved scaffold would not deviate from that of the experimentally X-ray resolved one. The present study investigates the suitability of alchemical free energy (AFE) methods for improving on this multistep docking and scoring protocol by means of a further re-scoring of ligands. AFE methods are increasingly used for predictions of free energies of binding in blinded competitions such as SAMPL (Statistic Assessment of Modelling of Proteins and Ligands) and D3R grand challenges [9–15]. Some AFE protocols have even achieved predictions of binding energies with root mean square deviations (RMSD) under 1.5 kcal mol^-1^, and Pearson Correlation coefficients (R) of around 0.7 or better [16–23]. Nevertheless, the performance varies significantly between different AFE protocols and targets [24–26] and it is important to explore further the robustness of these methodologies.

Specifically, this study aimed to explore the extent to which a setup protocol motivated by previous domain knowledge may influence the accuracy of AFE calculations, and whether issues such as binding poses selection or binding site water placement can be overcome via an increase of the simulation time. This was investigated using a dataset of 15 congeneric inhibitors of the protein activated Cdc42-associated kinase (ACK1) [27], a potential cancer target [28, 29]. The compounds span a large range of activity (K_i_ values ranging from more than 10 μM to 0.0002 μM), as seen in Table 1, and are typical of the structural modifications performed in hit-to-lead programs. The 15 ligands were first docked into the ACK1 ATP-binding site, and a set of docked poses obtained for each ligand was re-scored with a 4-step minimization protocol followed by a single-snapshot MM/PBSA re-scoring. The best scored pose was alchemically studied and the relative binding energy was compared to the experimental one. The alchemical calculations were also repeated with a 10-fold increase in sampling time. The role of a possible bridging water molecule in the binding pocket was also considered. Finally, thermodynamic cycle closures were analyzed as a way to detect incorrectly predicted poses without knowledge of the experimental relative binding energies.

**Table 1.** Ligands studied in this work, along with reported *K*_i_ values [27]. Compound numbering is the same as that used in reference 27.

| | | A | B | C | D | E | F | G | J | $K_i$ | Compound |
| --- | --- | --- | --- | --- | --- | --- | --- | --- | --- | --- | --- |
|  | Batch 1 | O | C |  | N | C | C | O | H | 0.01 | **2** |
|  |  | O | C |  | N | C | C | S | H | > 10 | **3** |
|  |  | O | C |  | N | C | N | N | H | 7.3 | **4** |
|  |  | O | C |  | N | N | C | N | H | 1.8 | **5** |
|  |  | O | C |  | C | C | C | O | H | 0.07 | **6** |
|  |  | O | C |  | N | C | C | NH | H | 0.006 | **7** |
|  | Linker | O | C |  | N | C | C | O | OCH<sub>3</sub> | 0.005 | **8** |
|  | Batch 2 | O | C |  | N | C | C | O | O(CH<sub>2</sub>)<sub>2</sub>NMe<sub>2</sub> | 0.005 | **15** |
|  |  | O | C |  | N | C | C | NH | O(CH<sub>2</sub>)<sub>2</sub>NMe<sub>2</sub> | 0.006 | **16** |
|  |  | S | S |  | N | C | C | NH | O(CH<sub>2</sub>)<sub>2</sub>NMe<sub>2</sub> | 0.0003 | **35** |
|  |  | S | S |  | N | C | C | O | O(CH<sub>2</sub>)<sub>2</sub>NMe<sub>2</sub> | 0.0002 | **36** |
|  |  | S | C |  | N | C | C | O | O(CH<sub>2</sub>)<sub>2</sub>NMe<sub>2</sub> | 0.08 | **38** |
|  |  | O | O |  | N | C | C | O | O(CH<sub>2</sub>)<sub>2</sub>NMe<sub>2</sub> | 0.013 | **39** |
|  |  | C | C | OH | N | C | C | NH | O(CH<sub>2</sub>)<sub>2</sub>NMe<sub>2</sub> | 0.04 | **44** |
|  |  | C | C | OMe | N | C | C | NH | O(CH<sub>2</sub>)<sub>2</sub>NMe<sub>2</sub> | 0.05 | **45** |

## MATERIALS AND METHODS

### Dataset

The dataset consists of 15 ACK1 competitive inhibitors for which inhibition constants (*K*_i_) have been reported. The structure of only one protein-ligand complex (compound **35**) was determined by X-ray crystallography [27] (Fig 1). This dataset was further divided into two subsets: *batch 1* (6 ligands with *K*_i_ values ranging from >10 μM to 0.006 μM), and *batch 2* (9 ligands with *K*_i_ values ranging from 0.013 to 0.0002 μM).

**Fig 1.**
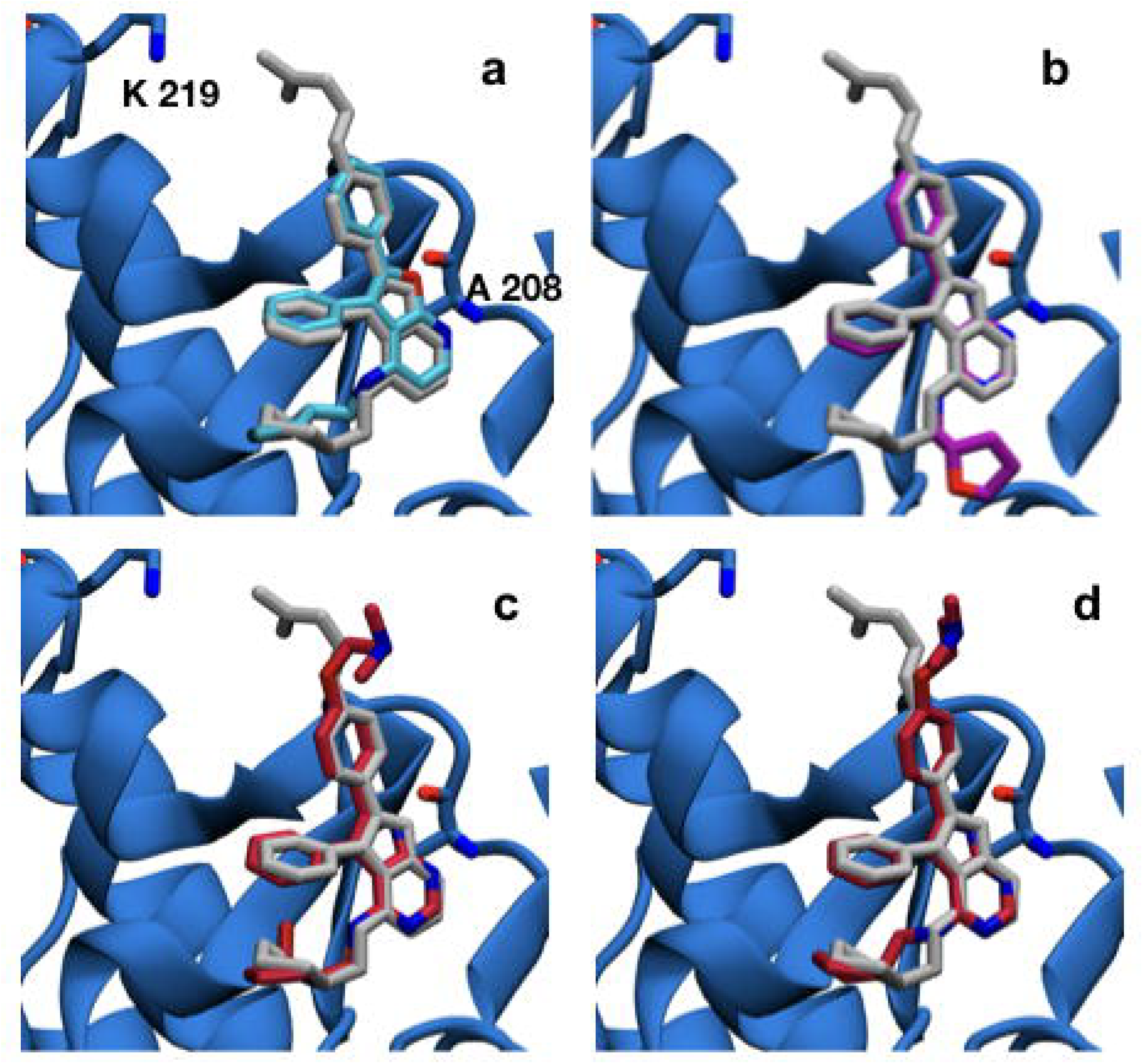
(a-d) Superimposition of the X-ray diffraction derived structure of the ACK1 protein co-crystallized with ligand 35 (grey) (PDB code 4EWH). (a) with the best predicted MM/PBSA docked pose for ligand **6** (blue), (b) with ligand 7 (purple) exhibiting a different binding mode, (c) with ligand **44** from MM/PBSA prediction using the best predicted binding mode and (d) with ligand **44** using the second-best binding mode prediction. All carbon atoms of ligand **44** are colored in red. Hydrogens are omitted for clarity.

### Protein and ligand setup

The ACK1 kinase domain structure was taken from the Protein Data Bank, code 4EWH [27], using chain B of the crystal structure, which was protonated with MOE v2009.1 [30]. The structure has no missing residues; Tyrosine 284 was dephosphorylated with MOE following Lougheed *et al.* observation that inhibitor binding is not expected to be sensitive to the phosphorylation state of this residue [31]. The protonation state of each ligand was predicted using the SDwash program in MOE v2009.1. *Batch 1* ligands were predicted to be neutral, whereas *batch 2* ligands were predicted to be positively charged.

### Docking

Docking was performed with MOE v2009.1 [30]. The full docking process was done in three steps. The first one was an exhaustive conformational search of the ligands using the *Systematic* option of MOE together with the option *Enforce chair conformations* on. All other parameters were set to the standard options. A maximum of 100 conformations by compound were selected for the *Placement* step. In the second step the receptor was defined as those atoms within 9.0 Å from the ligand. The *Rotate Bonds* option was activated and the *Affinity dG function* employed together with the *Triangle Matcher* method for placement. A maximum of 30 poses for each ligand were retained. Finally, the 500 best structures were submitted to the *Refinement* step with the *Force Field* function and allowing the lateral chains of the pocket residues to move during the optimization without restriction. All other parameters were set to the standard options. The five best structures obtained for each ligand, according to their predicted binding energies, were retained for minimization and re-scoring with MM/PBSA.

### MM/PBSA

A four-step minimization protocol followed by a single snapshot MM/PBSA re-scoring was performed with Amber 14 [32]. Ligands were prepared with Antechamber using the GAFF force field [33] with AM1-BCC partial charges [34, 35], while the ff99SB [36] force field was used for the protein. All systems were solvated in a rectangular box of TIP3P water molecules [37]. Counterions were added as necessary to neutralize the systems [38]. Energy minimization was performed under periodic boundary conditions using the particle-mesh-Ewald method for the treatment of the long-range electrostatic interactions [39]. A cut-off distance of 10 Å was chosen to compute non-bonded interactions. The four-step minimization procedure was as follows: 1) 5000 steepest descent (SD) steps applied to water molecule coordinates only; 2) 5000 SD steps applied also to protein atoms, with positional harmonic restraints (5 kcal mol^-1^ Å^-2^) applied to backbone atoms only; 3) 5000 SD steps as done previously with backbone atom restraints set to 1 kcal mol^-1^ Å^-2^ and 4) 5000 SD steps with no restraints.

For each of the energy minimized structures, a binding free energy was estimated following the MM/PBSA method using the MM/PBSA.py program [40]. No entropic contributions were taken into account, while the variables *cavity_surften* and *cavity_offset* were assigned the values of 0.00542 kcal mol^-2^ Å^-2^ and −1.008, respectively, using the defaults for all remaining variables.

### Alchemical free energy calculations

Relative binding free energies were calculated using a single topology molecular dynamics alchemical free energy approach [41]. Alchemical free energy calculations avoid direct computation of the free energy change associated with the reversible binding of a ligand to a protein through an artificial morphing of a ligand *X* into another ligand *Y* by using a parameter *λ* which defines the change from *X* to *Y*. Thus, the relative free energy of binding (ΔΔ*G*_*X→Y*_) was given by equation 1 as:

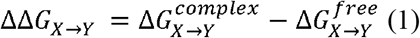

Where 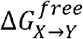 is the free energy change for transforming ligand *X* into ligand *Y* in solution whereas 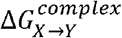 is the free energy change for the same transformation in the protein binding site. A relative free energy perturbation network for both *batch 1* and *batch 2* was designed (S5 Fig and S14 Fig). The top-scored MM/PBSA pose for each ACK1 ligand was used as input for the subsequent alchemical free energy preparation protocol using the FESetup software package [42]. The protocol used by FESetup for the automated preparation of ligands, protein and complexes was as follows:

#### Ligands

Atomic charges were assigned by using the Antechamber module in AmberTools 14 [32], selecting the AM1-BCC method [34, 35], and the GAFF2 force field [33]. Ligands were solvated with TIP3P water molecules [37], with counterions added as necessary to neutralize the system [38]. Each system was energy minimized for 100 SD cycles and equilibrated at 300 K and 1 atm pressure for 10^5^ molecular dynamics (MD) steps with a 2 fs timestep using the module Sander [32], with a positional harmonic restraint (10 kcal mol^-1^ Å^-2^) applied to ligand atoms. Bonds involving hydrogen atoms were constrained.

#### Protein

The protein was parametrized using the Amber ff14SB force field [43].

#### Complexes

Each ligand was combined back with the ACK1 protein model and the complex was solvated with TIP3P water molecules [37]. Counterions were also added to neutralize the solution [38]. The system was afterwards equilibrated following the procedure already described for ligands, using now 5000 MD steps.

All alchemical free energy calculations used 11 equidistant λ windows. For each λ value MD trajectories were computed in the NPT ensemble with a pressure of 1 atm and temperature of 300 K using the software SOMD 2016.1.0 [25, 44]. SOMD has been used in several recent studies to model the binding energetics of enzyme inhibitors [26], carbohydrate ligands [24], and host-guest systems [13]. Each λ window was sampled for 4 ns. Pressure was regulated using a Monte Carlo barostat [45, 46] with an update frequency of 25 MD steps. Temperature was kept constant using the Andersen thermostat [47], with a collision frequency of 10 ps^-1^. A 2 fs time step was used with the leapfrog-Verlet integrator. All bonds involving hydrogens were constrained to their equilibrium distances. Non-bonded interactions were evaluated setting a cut-off distance of 12 Å. Long-range electrostatic interactions were calculated using the shifted atom-based Barker-Watts reaction field [48], with the medium dielectric constant set to 82.0. In order to avoid steric clashes at the beginning of each MD run due to modifications of the ligand parameters associated with changes in λ, each structure was energy minimized for 1000 steps prior to MD simulation.

Each simulation was repeated at least twice using different initial assignments of velocities, and both ΔΔG_X→Y_ and ΔΔG_Y→X_ were calculated from independent simulations. In some cases, when poor agreement was observed between duplicates a third run was performed.

Ligand **38** was tested as a racemic mixture for consistency with the experimental conditions. Calculations were carried out for each enantiomer and the binding energies relative to this ligand were given with equation 2:

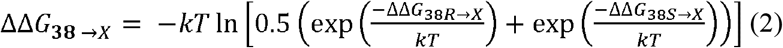

Cycle closures were evaluated using free energy changes from both the forward (**X→Y**) and reverse (**Y→X**) perturbations. The metrics used to evaluate the datasets were the determination coefficient R^2^, linear regression slope and the mean unsigned error (MUE). Experimental binding affinities were calculated from the corresponding inhibition constants [27] (K_i_) using 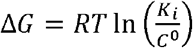 with C^0^ = 1 mol L^-1^. As no uncertainties have been reported for the *K_i_* values, an uncertainty of 0.4 kcal mol^-1^ was assumed [20, 49].

Relative free energies were estimated using the multistate Bennett’s acceptance ratio (MBAR) method [50], as included in the software *analyse_freenrg* from the Sire software suite. Relative free energies for complete datasets were evaluated using version 0.3.5 of the *freenrgworkflows* python module [https://github.com/michellab/freenrgworkflows], using ligand **3** as a reference, for which a K_i_ = 10 μM is used.

For more details, see Mey *et al.* [51]. All analysis scripts are available online at https://github.com/michellab/ACK1_Data.

#### Alchemical free energy Protocols

Five different alchemical free energy protocols were followed. *Protocol A* uses for each ligand the best scored pose according to MM/PBSA. This leads to a pose that differs from the X-ray crystallographic pose of **35** for several ligands (**2, 4, 7, 8, 16, 44** and **45**). *Protocol B* needs user intervention to select the pose most similar to the experimental binding mode. *Protocols C* and *D* explore the effect of manually modelling a water molecule inside the ACK1 ATP-binding site (see Fig 2). This reflects user knowledge that in other high-resolution structures of ACK1 (e.g. the 1.31 Å resolution 4HZR structure [52]) one additional binding site water molecule between the protein and ligand is apparent. *Protocol C* uses the same ligand poses as *Protocol A*, while *Protocol D* uses the same poses as *Protocol B*.

**Fig 2.**
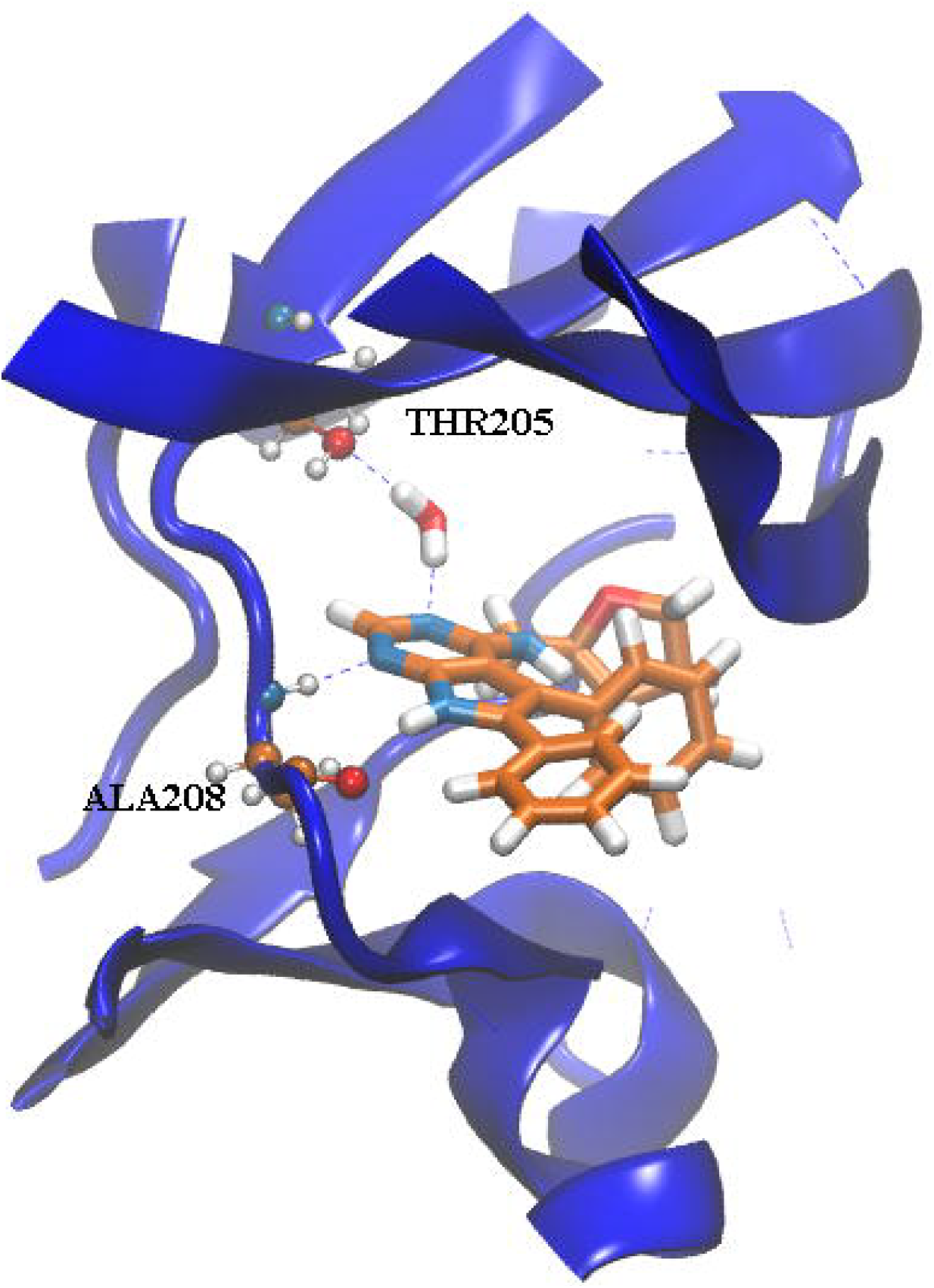
Snapshot taken after 2 ns of MD for the λ=0, run 1, 7→6 perturbation, showing the manually placed water molecule inside the ATP-binding pocket.

Finally, *Protocol E* is simply *Protocol A* with the per *λ* simulation time increased ten-fold. This was done to evaluate whether the different binding mode and ATP-binding site water rearrangements seen in *Protocols A-D* can be sampled with longer MD simulation protocols. *Protocol E* is computationally expensive and was applied to *batch 1* only (ca. 10 μs of simulation time).

The stability of the ligand poses and protein structure for all protocols was assessed by plotting the distribution of RMSD values across the dataset for the ligand and the protein backbone atoms (S1 and S2 Figs). The results indicate that the poses are generally stable (RMSD < 2 Å for almost all poses), the protein structure is generally stable (mean RMSD ca. 1.5 Å), though in some instances large positional fluctuations of the distal N-terminal region contribute to an increased RMSD.

Figures were rendered with VMD [53], while graphs were prepared with Origin [54] and python using plotting libraries Matplotlib version 2.0.2 [55] and Seaborn version 0.7.1 [56].

## RESULTS

### Batch 1

*Protocol A* renders (Table 2) modest results, with a R^2^ of 0.36±0.07 and a strong underestimation of relative free energies, as shown by the slope of the regression line (0.3). Inspection of Fig 3a and S3 Fig shows that ligands **2, 4** and **7** are clear outliers. These ligands have a predicted docked pose which differ more from the X-ray derived binding mode (see Figs 1b and 1c). Results for *protocol B* are shown in Table 2, and S3, S6 and S9 Figs. This protocol gives clearly better results, although the underestimation (slope 0.4) of relative binding free energies remains high, and ligands **2, 4** and **7** are still ranked poorly.

**Table 2.** R^2^, MUE (kcal mol^-1^) and slope metrics obtained from the comparison of experimental ^*a*^ and predicted relative free energies of binding of *batch 1*.

| Protocol | $R^2$ | MUE | Slope |
| --- | --- | --- | --- |
| A | $0.36 \pm 0.07$ | $1.55 \pm 0.06$ | 0.3 |
| B | $0.67 \pm 0.03$ | $1.29 \pm 0.03$ | 0.4 |
| C | $0.5 \pm 0.2$ | $1.53 \pm 0.07$ | 0.2 |
| D | $0.84 \pm 0.03$ | $1.29 \pm 0.03$ | 0.5 |
| E | $0.33 \pm 0.08$ | $1.73 \pm 0.08$ | 0.3 |

<sup>a</sup> An upper bound of $R^2 = 0.97 \pm 0.02$ on achievable predictions may be estimated given assumed experimental uncertainties of 0.4 kcal.mol<sup>-1</sup>

**Fig 3.**
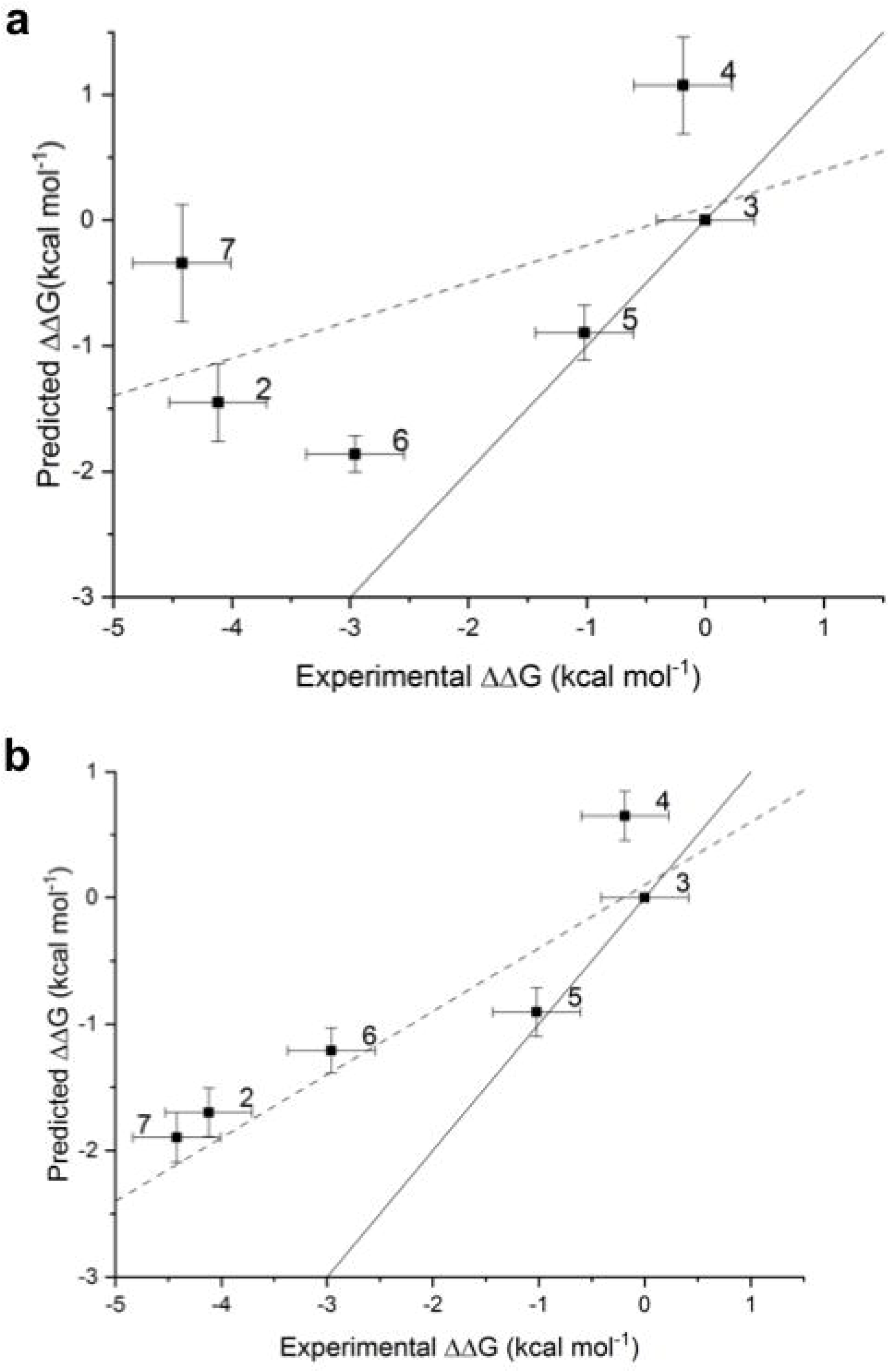
(a) and (b) Comparison of experimental and predicted relative free energies of binding of *batch 1* for *protocols A* and *D*, respectively. Free energies of binding are relative to ligand **3**. The linear regression line (dashed line) and a line with slope unity (solid line) are also presented.

An analysis of the relative binding energies calculated with *protocols A* and *B* (S3, S5 and S6 Figs), for ligands **2, 4** and **7**, reveals that these ligands appear in the perturbations that show the highest deviations between the experimental and calculated relative binding energies. Thus, for *protocol A* deviations of more than 3.0 kcal mol^-1^ are observed for **2→3, 7→4, 7→6, 7→3** and **3→7**, while for *protocol B* these deviations appear for perturbations **2→3, 4→7, 7→4, 7→3** and **3→7**. An analysis of the docked structures of ligands **6** and **7** suggested that a possible explanation for the inability of the protocol to reproduce the experimental relative binding affinities is due to interactions of the extra nitrogen atom in the pyrimidine ring of ligand **7** that is missing in the pyridine ring of ligand **6** (see Fig 4). The extra N atom in the pyrimidine ring could establish a hydrogen bond with THR205 (see Fig 2) if a bridging water was present. Indeed, several water molecules are present inside the ATP-binding pocket of 4HZR [52]. That possibility was explored in *protocols C* and *D*, where a water molecule was manually placed inside the binding pocket between the nitrogen in position D of ligand **7** (see Table 1) and THR205. The final position of the water molecule is obtained after 100 steps of SD minimization fixing all other atoms. Results for *protocol C* are shown in Table 2 and S10 Fig, while those for *protocol D* appear in Table 2 and Figs 3b and 4. *Protocol D* clearly surpass all others, with a R^2^ of 0.84±0.03 and an improvement in the underestimation of relative binding energies (slope = 0.5). A comparison of the calculated relative binding energies for ligands **3** and **4** allows to conclude that using a different pose for ligand **4** does not seem to affect the results (both *protocols A* and *B* for example, give an average ΔΔG**_3→4_** of 1.3 kcal mol^-1^). Inspection of the calculated trajectories show that ligand **4** rapidly converts from its initial docked pose (*protocols A* and *C*) to one similar to that used as input for *protocols B* and *D*. The computed trajectories were visualized to assess the stability of the active site water molecule. The water molecule showed little drift from its initial position in most cases, with the exception of perturbations involving compound 6 in protocol C, where the water frequently diffused away from the binding site.

**Fig 4.**
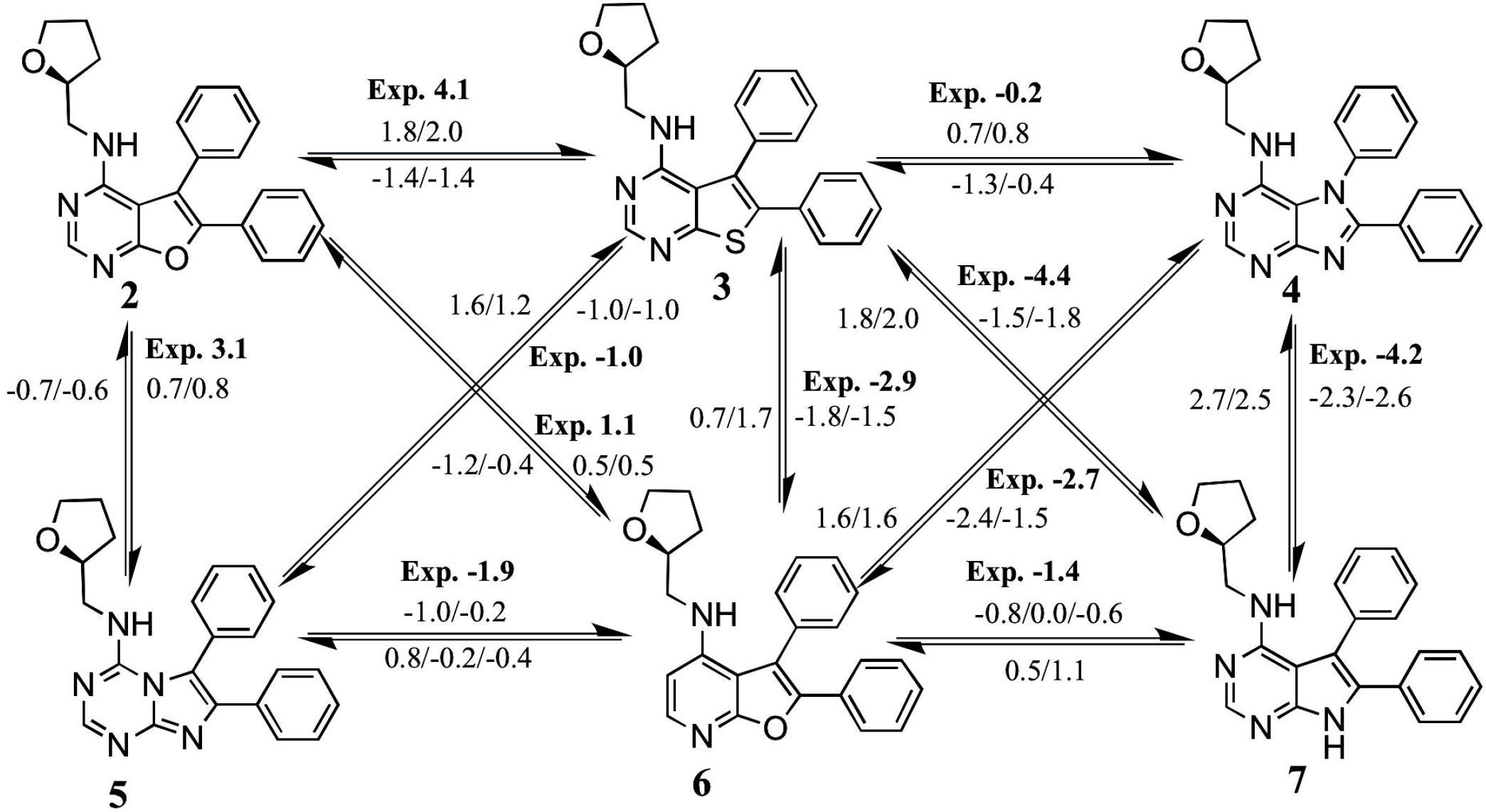
Calculated and experimental (in bold) relative binding affinities (in kcal mol^-1^) for all the perturbations run in *batch 1* with *protocol D*. The calculated values correspond to independent repeats.

The possibility of resolving ambiguities in binding poses and binding site water content without user intervention was next tested by increasing the simulation sampling time to 40 ns for each λ window. The expectation was this would allow the ligand to find the correct pose and to allow water molecules diffuse in the ATP-binding site (see Fig 2). Results are shown in Table 2 and S3 and S11 Figs. The increased simulation time does not translate into any improvement of the results. The R^2^, slope and MUE values are as poor or poorer as those for *protocol A*, while the outliers remain the same. The MD trajectories show that, even with the increased simulation time, ligand **7** is not able to change its docking pose, while ligand **4** needs under 4 ns to adopt a pose that resembles the X-ray pose of **35**. Besides, a water molecule enters and remains in the ATP-binding site in 7 out of 22 MD trajectories only.

### Analysis of the complete dataset

The robustness of the results obtained for *batch 1* was tested by processing *batch 2* and re-analyzing the full dataset. Ligands in *batch 2* are positively charged in the assay conditions, whereas *batch 1* ligands are neutral. Relative free energy calculations that involve a net charge change are still technically challenging for simulations carried out with a reaction-field cut-off. Thus, the perturbations between ligands **8** and **15** were carried out assuming **15** is neutral. This is of course a significant simplification. Results for individual perturbations in *batch 2* are shown in S4 and S14 to S18 Figs.

*Protocol A*, as expected given the results obtained for *batch 1*, gives modest results, as can be seen in Table 3 and Fig 5a (R^2^ =0.45±0.06 and slope of 0.5). The slope has improved from 0.3 to 0.5 because the relative free energies of compounds in *batch 2* are not as under predicted as those from *batch 1* (see S1 Table). Ligands **16, 44** and **45** need further inspection. S14 Fig shows that, while the experimental ΔΔG**_45→44_** is −0.1 kcal mol^-1^, the calculated ΔΔG**_45→44_** are 1.8/1.5 (run 1/run 2) kcal mol^-1^ (the reverse perturbation was calculated as −2.0/-1.9 kcal mol^-1^). Similarly, while the experimental ΔΔG**_16→45_** is 1.2 kcal mol^-1^, the calculated results are ΔΔG**_16→45_** −0.8/-1.6 (run 1/run 2) kcal mol^-1^ and ΔΔG**_45→16_** −2.2/-2.4 /run 1/run 2) kcal mol^-1^.

**Table 3.**
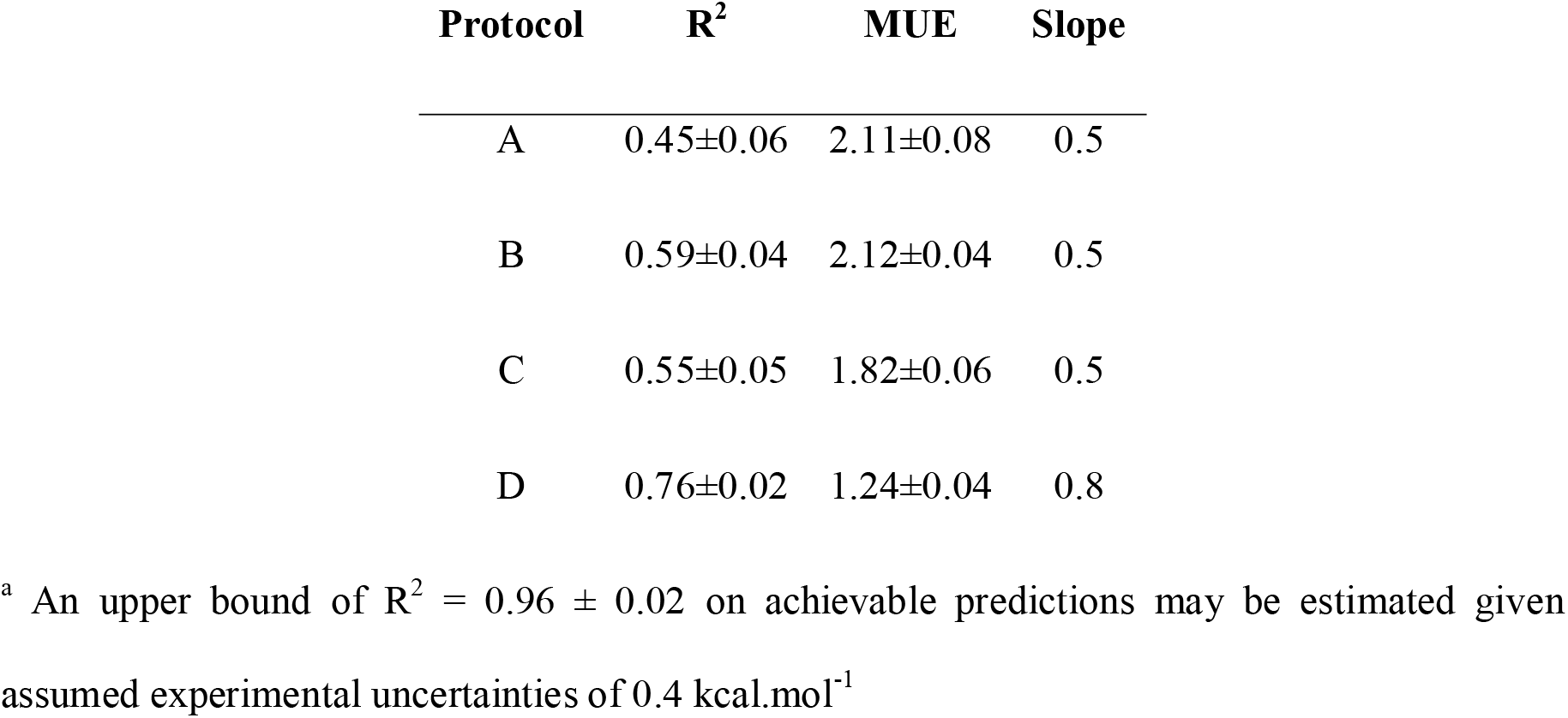
R^2^, MUE (kcal mol^-1^) and slope metrics obtained from the comparison of experimental^a^ and predicted relative free energies of binding of the whole set.

| Protocol | $R^2$ | MUE | Slope |
| --- | --- | --- | --- |
| A | 0.45±0.06 | 2.11±0.08 | 0.5 |
| B | 0.59±0.04 | 2.12±0.04 | 0.5 |
| C | 0.55±0.05 | 1.82±0.06 | 0.5 |
| D | 0.76±0.02 | 1.24±0.04 | 0.8 |

<sup>a</sup> An upper bound of $R^2 = 0.96 \pm 0.02$ on achievable predictions may be estimated given assumed experimental uncertainties of 0.4 kcal.mol<sup>-1</sup>

**Fig 5.**
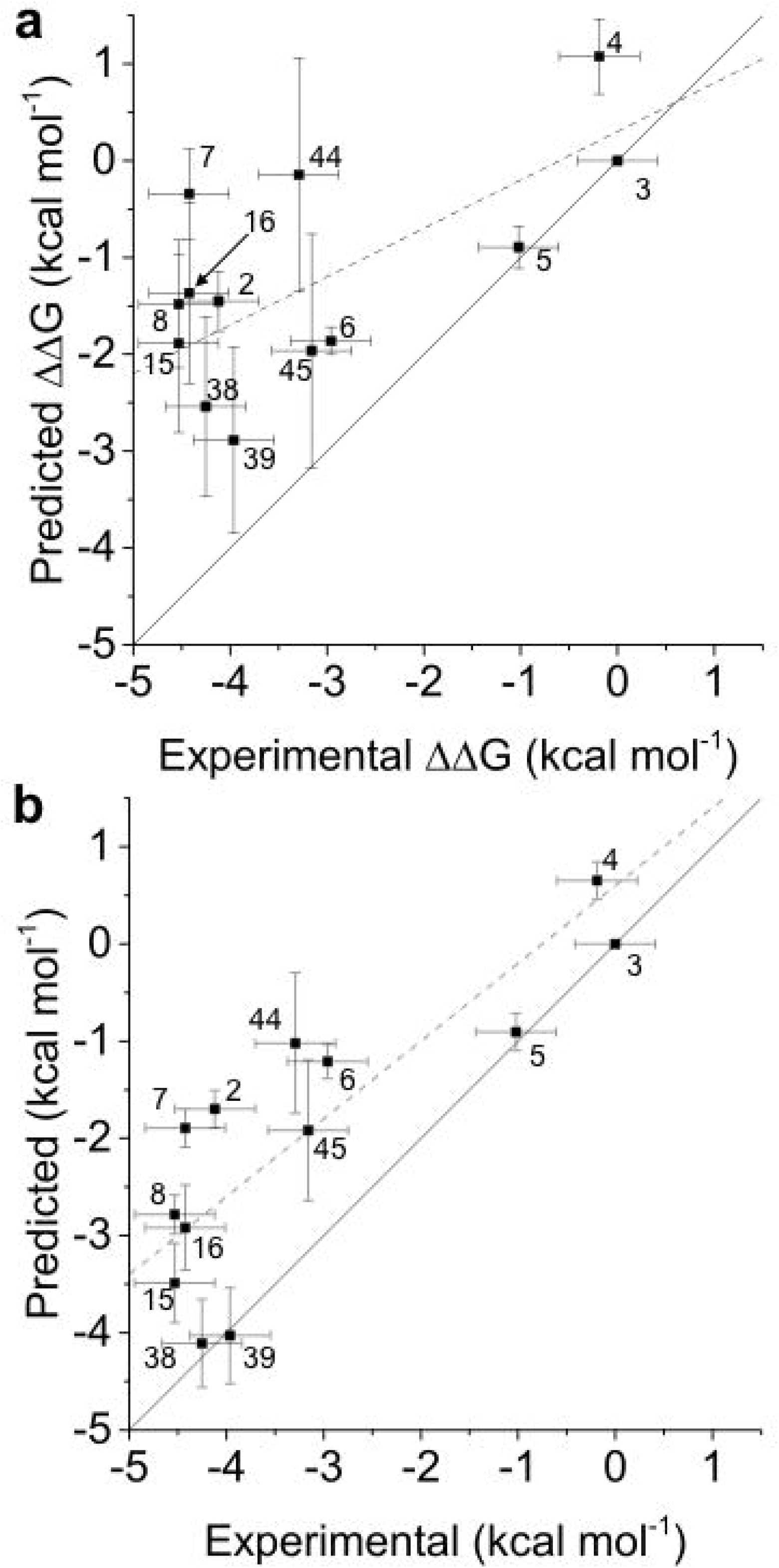
(a) and (b) Comparison of experimental and predicted relative free energies of binding of the whole set for *protocols A* and *D*, respectively. Free energies of binding are relative to ligand **3**. The linear regression line (dashed line) and a line with slope 1 (solid line) are also presented.

Interestingly, the dihedral angle defining the relative orientation of the NH group that links the pyrimidine and the cyclopentanol rings changes values rather quickly during the simulation. Fig 6a shows an example for the first repeat of the perturbation **44→45** at λ=0. For the simulations involving ligand **44** an intramolecular H-bond between its aniline NH group and its cyclopentyl hydroxyl group is established (see Fig 6c). That conformation is precisely the second-best MM/PBSA docked one (see Fig 1c), which features that intramolecular hydrogen bond. Thus, *batch 2 protocol B* includes the second-best scored MM/PBSA poses for ligands **8, 16** and **44**. In the case of ligands **8** and **16**, this implies using a pose that resembles the most the X-ray binding mode, while for ligand **44** the second-best scored MM/PBSA pose differs from the best-scored one in the aniline NH dihedral angle (see Fig 1c). The improvement, as shown in Table 3 and S12 Fig, for *protocol B* as compared with *protocol A*, is quite modest. Results are clearly better for the **16→45** and **45→16** perturbations, with the disagreement between experimental and calculated relative binding energy decreasing from 3.5 to 0.3 kcal mol^-1^ (compare S14 and S15 Figs), but ligand **44** is still an outlier. Although the experimental relative binding energy for the **45 → 44** perturbation is just −0.1 kcal mol^-1^, ligand **45** is predicted to bind much more strongly to ACK1 (calculated ΔΔG_**45**→ **44**_ are −2.0/−1.9 and −1.2/−1.6 kcal mol^-1^ for *protocols A* and *B*, respectively) than **44**. This suggests possible deficiencies in the force field used for **44** in this study.

**Fig 6.**
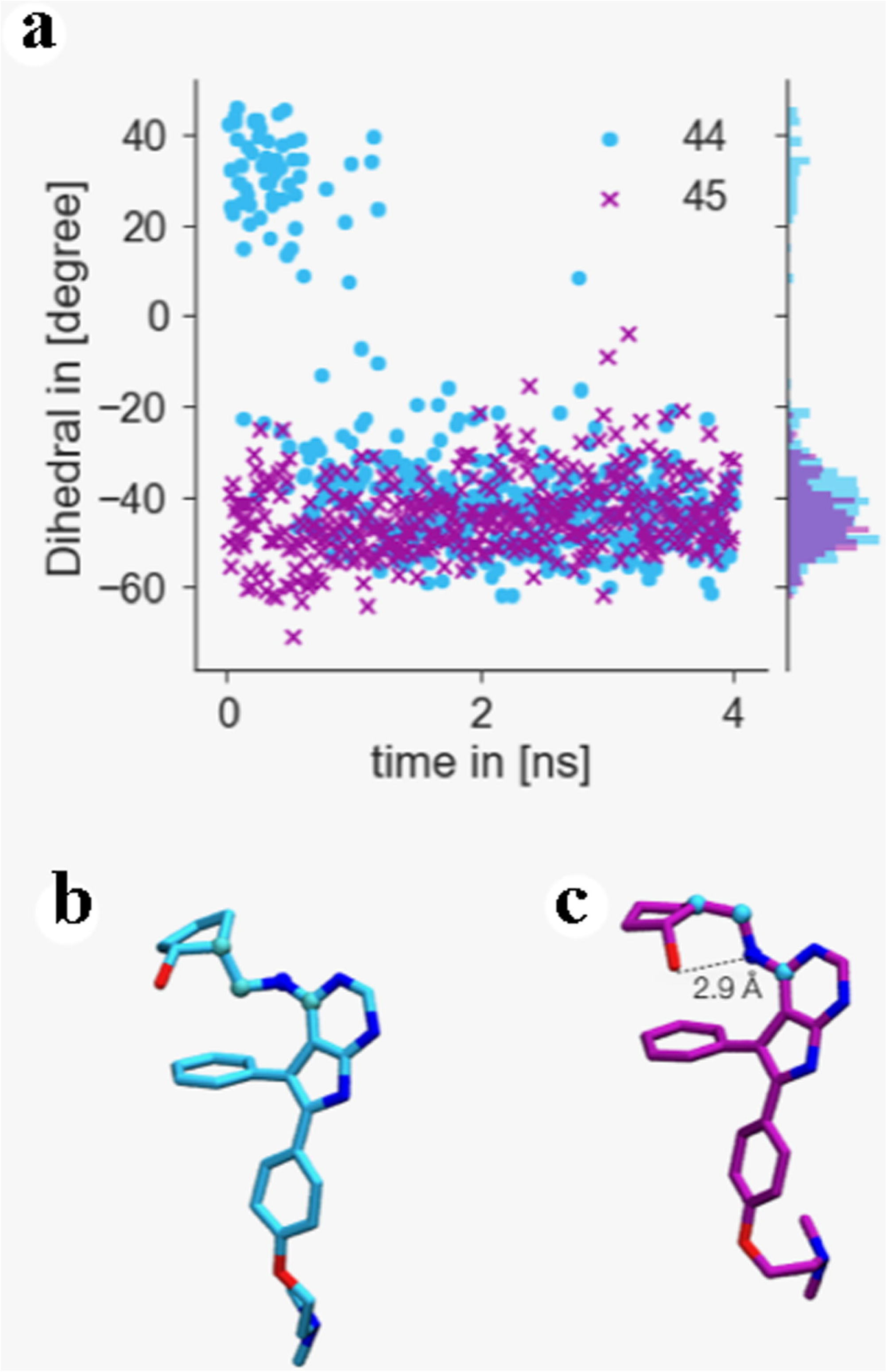
(a) 4 ns trajectory monitoring dihedral angle of ligand 44 (blue circles) and 45 (purple crosses) as indicated in (b) and (c) as well as probability distribution of dihedrals over the trajectory. (b) Snapshot of the conformation of ligand 44 taken from a λ = 0 trajectory at t=0 ns indicating dihedral conformation monitored in (a) highlighted by spheres. (c) Snapshot of the conformation of ligand 44 taken from the same trajectory after 3 ns, showing an intramolecular hydrogen bond.

*Protocols C* and D, follow the same trends already explained for *batch 1*, pointing to an improvement in the results when a water molecule is included in the ATP-binding pocket (Table 3). An encouraging R^2^ of 0.76 ± 0.02 and an improvement in the underestimation of relative binding energies (slope 0.8) is obtained, though there is still room for improvements for affinity predictions for **44** and **16**.

### Thermodynamic cycle closures analysis

Hysteresis, being defined as the difference in binding energy between the forward and reverse perturbation [44, 57, 58], has been proposed as useful metric to identify problematic perturbations [59, 60]. Cycle closures for both *batch 1* and *batch 2* were computed to determine whether incorrectly predicted binding poses could be detected in the absence of experimental binding affinity data. Results are shown in Table 4.

**Table 4.**
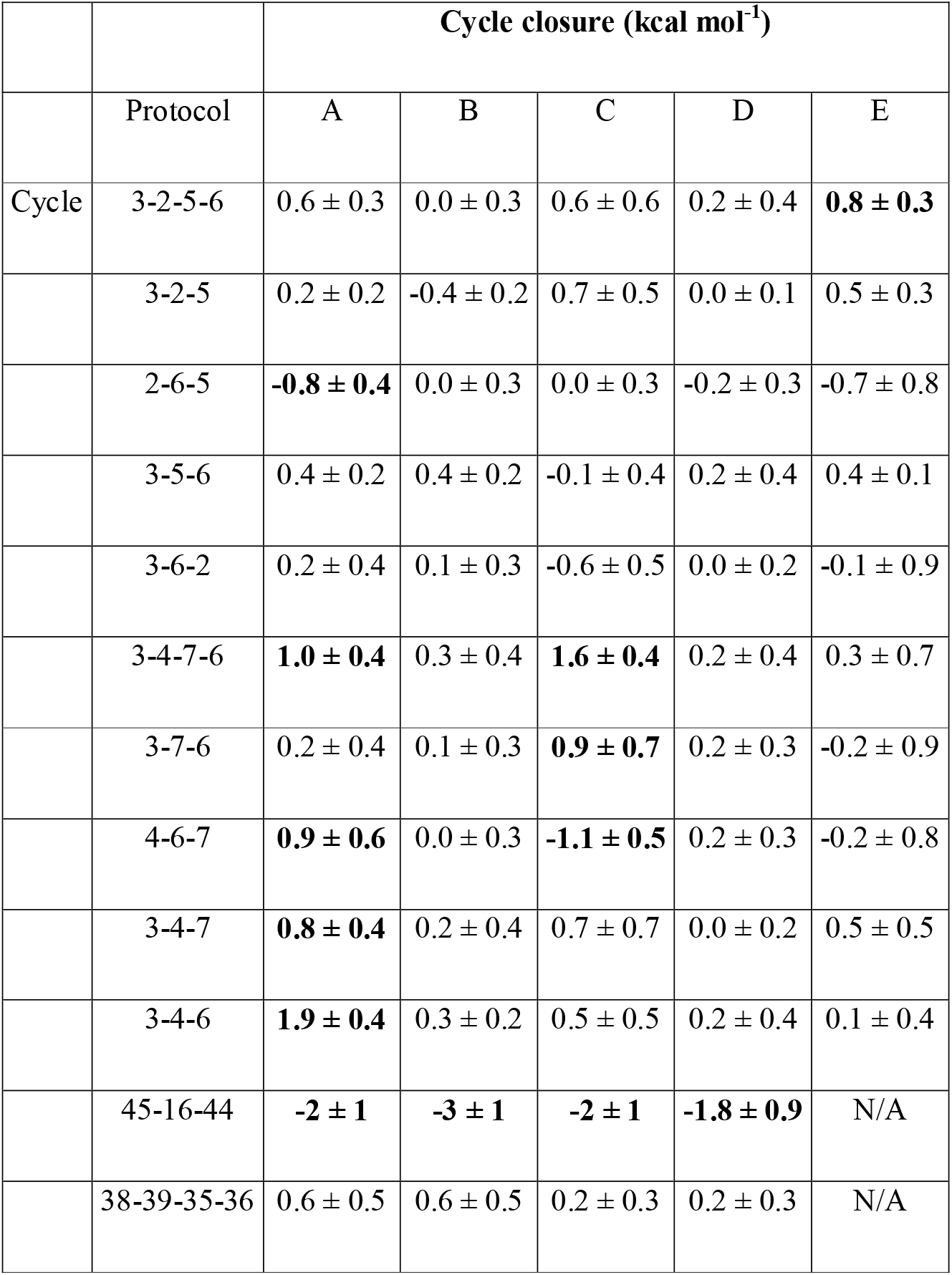
Calculated thermodynamic cycle closures. Cycle closures that exceed or equal a threshold of 0.8 kcal mol^-1^ are highlighted in bold.

|  | Protocol | A | B | C | D | E |
| --- | --- | --- | --- | --- | --- | --- |
| Cycle | 3-2-5-6 | $0.6 \pm 0.3$ | $0.0 \pm 0.3$ | $0.6 \pm 0.6$ | $0.2 \pm 0.4$ | **$0.8 \pm 0.3$** |
| | 3-2-5 | $0.2 \pm 0.2$ | $-0.4 \pm 0.2$ | $0.7 \pm 0.5$ | $0.0 \pm 0.1$ | $0.5 \pm 0.3$ |
| | 2-6-5 | **$-0.8 \pm 0.4$** | $0.0 \pm 0.3$ | $0.0 \pm 0.3$ | $-0.2 \pm 0.3$ | $-0.7 \pm 0.8$ |
| | 3-5-6 | $0.4 \pm 0.2$ | $0.4 \pm 0.2$ | $-0.1 \pm 0.4$ | $0.2 \pm 0.4$ | $0.4 \pm 0.1$ |
| | 3-6-2 | $0.2 \pm 0.4$ | $0.1 \pm 0.3$ | $-0.6 \pm 0.5$ | $0.0 \pm 0.2$ | $-0.1 \pm 0.9$ |
| | 3-4-7-6 | **$1.0 \pm 0.4$** | $0.3 \pm 0.4$ | **$1.6 \pm 0.4$** | $0.2 \pm 0.4$ | $0.3 \pm 0.7$ |
| | 3-7-6 | $0.2 \pm 0.4$ | $0.1 \pm 0.3$ | **$0.9 \pm 0.7$** | $0.2 \pm 0.3$ | $-0.2 \pm 0.9$ |
| | 4-6-7 | **$0.9 \pm 0.6$** | $0.0 \pm 0.3$ | **$-1.1 \pm 0.5$** | $0.2 \pm 0.3$ | $-0.2 \pm 0.8$ |
| | 3-4-7 | **$0.8 \pm 0.4$** | $0.2 \pm 0.4$ | $0.7 \pm 0.7$ | $0.0 \pm 0.2$ | $0.5 \pm 0.5$ |
| | 3-4-6 | **$1.9 \pm 0.4$** | $0.3 \pm 0.2$ | $0.5 \pm 0.5$ | $0.2 \pm 0.4$ | $0.1 \pm 0.4$ |
|  | 45-16-44 | **$-2 \pm 1$** | **$-3 \pm 1$** | **$-2 \pm 1$** | **$-1.8 \pm 0.9$** | N/A |
| | 38-39-35-36 | $0.6 \pm 0.5$ | $0.6 \pm 0.5$ | $0.2 \pm 0.3$ | $0.2 \pm 0.3$ | N/A |

As could be expected, similar conclusions can be obtained when analyzing ring cycle closures or comparing forward and reverse perturbations, although there are some cases with high deviations between experimental and calculated relative binding energies, while exhibiting almost null hysteresis for the forward and reverse perturbations (i.e. the perturbations between ligands **2** and **5** in *batch 1* and those between ligands **44** and **45** in *batch 2*).

Overall it appears that a threshold of ± 0.8 kcal mol^-1^ for cycle closure errors is useful to flag poses that need further attention even without prior knowledge of the experimental binding affinities. Thus, for *protocol A*, **3**-**4**-**7**-**6, 3**-**4**-**6, 3**-**4**-**7, 4**-**6**-**7, 2**-**6**-**5** and **45**-**16**-**44** thermodynamic cycle closures are indicative of problematic ligands. According to this metric, a significant improvement when using *protocol B* (only one thermodynamic cycle closure above the threshold) is seen, while a comparison between *protocols A* (6 cycles with hysteresis above the threshold) and *C* (4 cycles) suggest a modest improvement. Results for *batch 2* clearly indicate that ligands **44, 45** and **16** (hysteresis of −2 ± 1 kcal mol^-1^ in their thermodynamic cycle for *protocol A*) are much more problematic than ligands **35, 36, 38** and **39** (hysteresis of 0.2 ± 0.3 kcal mol^-1^ for *protocol A*). The best performing *protocol D* is unable to improve the hysteresis for the **45**-**16**-**44** thermodynamic cycle.

## DISCUSSION

This work has explored the viability of using alchemical free energy methods as a final filter in a cascade of computational methods for hit-to-lead virtual screens in the context of a dataset of ACK1 inhibitors. The two major limitations of AFE methods are the quality of the potential energy function used, and the extent to which the configurational sampling performed has captured relevant protein-ligand conformations [59, 60] In principle sufficient long simulations will relax a protein-ligand complex to the ligand pose and protein conformation preferred by the force field used. However, because computing time is limited in practical scenarios, AFE simulations typically afford only a few ns per window, which can make the calculated binding affinities sensitive to the selection of the starting conformations. This work indicates that use of experimental data to bias the selection of poses and setup of binding site water content could lead to significant performance improvements. While the dataset studied here is small, the lessons from this study are likely applicable to other binding sites. Of course, as illustrated with ligand **4**, even in cases where the MD simulations relax a previously modelled binding pose to one that closely resembles a pose inferred from X-ray data, the free energy calculations may still fail to reproduce the experimental binding affinities.

Careful analysis of literature structural data [52, 61, 62] was key to identify a conserved hydration site that was not modelled in the prior docking calculations. This knowledge was important to realize upon inspection of putative poses for some ligands in *batch 1* the feasibility of a hydrogen bonding interaction via a bridging water molecule. Gratifyingly modelling of this hydration site leads to significant accuracy improvements for several perturbations.

In principle, assuming an accurate potential energy function, these sampling issues could be dealt with by simply increasing the sampling time of the MD simulations. For the present dataset, we find that a one order of magnitude increase in sampling time was insufficient to bring about improvements in binding poses accuracy and binding site water content. Thus, at present it seems wise to pay attention to the starting conditions of the free energy calculations to maximize cost effectiveness. Where experimental data is lacking, a number of molecular modelling protocols have been proposed to determine location and energetics of important binding site water molecules [63–72].

## SUPPORTING INFORMATION

**S1 Fig. Distribution of ligand RMSDs.**

**S2 Fig. Distribution of protein backbone RMSDs**

**S3 Fig. Calculated and experimental relative binding affinities for all protocols essayed in *batch 1***. The minus sign used to label run (-1), run (-2) and run (-3) indicates that, in order to easily compare forward and reverse perturbations, the sign of the relative binding affinity has been changed.

**S4 Fig. Calculated and experimental relative binding affinities for all protocols essayed in *batch 2***. The minus sign used to label run (-1), run (-2) and run (-3) indicates that, in order to easily compare forward and reverse perturbations, the sign of the relative binding affinity has been changed.

**S5 Fig. Calculated and experimental (in bold) relative binding affinities (in kcal mol^-1^) for all the perturbations essayed in *batch 1, protocol A***. The drawing tries to reflect the different conformations adopted by ligands **2, 4** and **7**. The calculated values correspond to independent repeats.

**S6 Fig. Calculated and experimental (in bold) relative binding affinities (in kcal mol^-1^) for all the perturbations essayed in *batch 1, protocol B***. The calculated values correspond to independent repeats.

**S7 Fig. Calculated and experimental (in bold) relative binding affinities (in kcal mol^-1^) for all the perturbations essayed in *batch 1, protocol C***. The drawing tries to reflect the different conformations adopted by ligands **2, 4** and **7**. The calculated values correspond to independent repeats.

**S8 Fig. Calculated and experimental (in bold) relative binding affinities (in kcal mol^-1^) for all the perturbations essayed in *batch 1*, protocol E.** The drawing tries to reflect the different conformations adopted by ligands **2, 4** and **7**. The calculated values correspond to independent repeats.

**S9 Fig. Comparison of experimental and predicted relative free energies of binding of *batch 1* for *protocol B***. Free energies of binding are relative to ligand **3**. The linear regression line (dashed line) and a line with slope 1 (solid line) are also presented.

**S10 Fig. Comparison of experimental and predicted relative free energies of binding of *batch 1* for *protocol C***. Free energies of binding are relative to ligand **3**. The linear regression line (dashed line) and a line with slope 1 (solid line) are also presented.

**S11 Fig. Comparison of experimental and predicted relative free energies of binding of *batch 1* for *protocol E***. Free energies of binding are relative to ligand **3**. The linear regression line (dashed line) and a line with slope 1 (solid line) are also presented.

**S12 Fig. Comparison of experimental and predicted relative free energies of binding of the whole set for *protocol B***. Free energies of binding are relative to ligand **3**. The linear regression line (dashed line) and a line with slope 1 (solid line) are also presented.

**S13 Fig. Comparison of experimental and predicted relative free energies of binding of the whole set for *protocol C***. Free energies of binding are relative to ligand **3**. The linear regression line (dashed line) and a line with slope 1 (solid line) are also presented.

**S14 Fig. Calculated and experimental (in bold) relative binding affinities (in kcal mol^-1^) for all the perturbations essayed in *batch 2, protocol A***. The drawing tries to reflect the different conformations adopted by ligands **44** and **16**. The calculated values correspond to independent repeats.

**S15 Fig. Calculated and experimental (in bold) relative binding affinities (in kcal mol^-1^) for all the perturbations essayed in *batch 2, protocol B***. The calculated values correspond to independent repeats.

**S16 Fig. Calculated and experimental (in bold) relative binding affinities (in kcal mol^-1^) for all the perturbations essayed in *batch 2, protocol C***. The drawing tries to reflect the different conformations adopted by ligands **44** and **16**. The calculated values correspond to independent repeats.

**S17 Fig. Calculated and experimental (in bold) relative binding affinities (in kcal mol^-1^) for all the perturbations essayed in *batch 2, protocol D***. The calculated values correspond to independent repeats.

**S18 Fig. Calculated and experimental (in bold) relative binding affinities (in kcal mol^-1^) for the perturbations used to link *batch 1* and *batch 2* with *protocols A, B, C* and *D***. The calculated values correspond to independent repeats.

**S1 Table. R^2^, MUE (kcal mol^-1^) and slope obtained from the comparison of experimental^a^ and predicted relative free energies of binding of *batch 2***

## REFERENCES

1. PhRMA. Fact Sheet “Drug Discovery and Development. Understanding theR&D process” 2017 [30.08.2017]. Available from: http://www.phrma.org/graphic/four-facts-about-spending-on-prescription-medicines.

2. Whitesides GM, Krishnamurthy VM. Designing ligands to bind proteins. Q Rev Biophys. 2006;38(4):385–95. Epub 07/03. doi:10.1017/S0033583506004240.

3. Smith AJT, Zhang X, Leach AG, Houk KN. Beyond Picomolar Affinities: Quantitative Aspects of Noncovalent and Covalent Binding of Drugs to Proteins. J Med Chem. 2009;52(2):225–33. doi:10.1021/jm800498e.

4. Kollman PA, Massova I, Reyes C, Kuhn B, Huo S, Chong L, et al. Calculating Structures and Free Energies of Complex Molecules:1 Combining Molecular Mechanics and Continuum Models. Acc Chem Res. 2000;33(12):889–97. doi:10.1021/ar000033j.

5. Aqvist J, Marelius J. The linear interaction energy method for predicting ligand binding free energies. Comb Chem High Throughput Screen. 2001;4(8):613–26. PubMed PMID: 11812258.

6. Zwanzig RW. High-Temperature Equation of State by a Perturbation Method. I. Nonpolar Gases. J Chem Phys. 1954;22(8):1420–6. doi:10.1063/1.1740409.

7. Michel J, Foloppe N, Essex JW. Rigorous Free Energy Calculations in Structure-Based Drug Design. Mol Inform. 2010;29(8-9):570–8. doi:10.1002/minf.201000051. PubMed PMID: 27463452.

8. Granadino-Roldan JM, Garzon A, Gomez-Gutierrez P, Pasamontes-Funez I, Santos Tomas M, Rubio-Martinez J. A multistep docking and scoring protocol for congeneric series: Implementation on kinase DFG-out type II inhibitors. Future Med Chem. 2018;10(3):297–318. doi:10.4155/fmc-2017-0156. PubMed PMID: 29338349.

9. Yin J, Henriksen NM, Slochower DR, Shirts MR, Chiu MW, Mobley DL, et al. Overview of the SAMPL5 host–guest challenge: Are we doing better? J Comput Aided Mol Des. 2017;31(1):1–19. doi:10.1007/s10822-016-9974-4.

10. Muddana HS, Fenley AT, Mobley DL, Gilson MK. The SAMPL4 host-guest blind prediction challenge: an overview. J Comput Aided Mol Des. 2014;28(4):305–17. doi:10.1007/s10822-014-9735-1. PubMed PMID: 24599514; PubMed Central PMCID: PMCPMC4053502.

11. Deng N, Flynn WF, Xia J, Vijayan RS, Zhang B, He P, et al. Large scale free energy calculations for blind predictions of protein-ligand binding: the D3R Grand Challenge 2015. J Comput Aided Mol Des. 2016;30(9):743–51. doi:10.1007/s10822-016-9952-x. PubMed PMID: 27562018.

12. Gaieb Z, Liu S, Gathiaka S, Chiu M, Yang H, Shao C, et al. D3R Grand Challenge 2: blind prediction of protein-ligand poses, affinity rankings, and relative binding free energies. J Comput Aided Mol Des. 2018;32(1):1–20. doi:10.1007/s10822-017-0088-4. PubMed PMID: 29204945; PubMed Central PMCID: PMCPMC5767524.

13. Bosisio S, Mey A, Michel J. Blinded predictions of host-guest standard free energies of binding in the SAMPL5 challenge. J Comput Aided Mol Des. 2017;31(1):61– 70. doi:10.1007/s10822-016-9933-0. PubMed PMID: 27503495.

14. Mey A, Juarez-Jimenez J, Hennessy A, Michel J. Blinded predictions of binding modes and energies of HSP90-alpha ligands for the 2015 D3R grand challenge. Bioorg Med Chem. 2016;24(20):4890–9. doi:10.1016/j.bmc.2016.07.044. PubMed PMID: 27485604.

15. Athanasiou C, Vasilakaki S, Dellis D, Cournia Z. Using physics-based pose predictions and free energy perturbation calculations to predict binding poses and relative binding affinities for FXR ligands in the D3R Grand Challenge 2. J Comput Aided Mol Des. 2018;32(1):21–44. doi:10.1007/s10822-017-0075-9. PubMed PMID: 29119352.

16. Wang L, Wu Y, Deng Y, Kim B, Pierce L, Krilov G, et al. Accurate and reliable prediction of relative ligand binding potency in prospective drug discovery by way of a modern free-energy calculation protocol and force field. J Am Chem Soc. 2015;137(7):2695–703. doi:10.1021/ja512751q. PubMed PMID: 25625324; PubMed Central PMCID: PMCUB153.

17. Shoichet BK, Walters WP, Jiang H, Bajorath J. Advances in Computational Medicinal Chemistry: A Reflection on the Evolution of the Field and Perspective Going Forward. J Med Chem. 2016;59(9):4033–4. doi:10.1021/acs.jmedchem.6b00511. PubMed PMID: 27054949.

18. Steinbrecher TB, Dahlgren M, Cappel D, Lin T, Wang LL, Krilov G, et al. Accurate Binding Free Energy Predictions in Fragment Optimization. J Chem Inf Mod. 2015;55(11):2411–20. doi:10.1021/acs.jcim.5b00538. PubMed PMID: WOS:000365465400011.

19. Homeyer N, Stoll F, Hillisch A, Gohlke H. Binding Free Energy Calculations for Lead Optimization: Assessment of Their Accuracy in an Industrial Drug Design Context. Journal of Chemical Theory and Computation. 2014;10(8):3331–44. doi:10.1021/ct5000296.

20. Mikulskis P, Genheden S, Ryde U. A Large-Scale Test of Free-Energy Simulation Estimates of Protein–Ligand Binding Affinities. J Chem Inf Mod. 2014;54(10):2794–806. doi:10.1021/ci5004027.

21. Ciordia M, Pérez-Benito L, Delgado F, Trabanco AA, Tresadern G. Application of Free Energy Perturbation for the Design of BACE1 Inhibitors. J Chem Inf Mod. 2016;56(9):1856–71. doi:10.1021/acs.jcim.6b00220.

22. Keränen H, Pérez-Benito L, Ciordia M, Delgado F, Steinbrecher TB, Oehlrich D, et al. Acylguanidine Beta Secretase 1 Inhibitors: A Combined Experimental and Free Energy Perturbation Study. Journal of Chemical Theory and Computation. 2017;13(3):1439–53. doi:10.1021/acs.jctc.6b01141.

23. Kuhn B, Tichý M, Wang L, Robinson S, Martin RE, Kuglstatter A, et al. Prospective Evaluation of Free Energy Calculations for the Prioritization of Cathepsin L Inhibitors. J Med Chem. 2017;60(6):2485–97. doi:10.1021/acs.jmedchem.6b01881.

24. Mishra SK, Calabró G, Loeffler HH, Michel J, Koča J. Evaluation of Selected Classical Force Fields for Alchemical Binding Free Energy Calculations of Protein-Carbohydrate Complexes. Journal of Chemical Theory and Computation. 2015;11(7):3333–45. doi:10.1021/acs.jctc.5b00159.

25. Mey ASJS, Juárez-Jiménez J, Hennessy A, Michel J. Blinded predictions of binding modes and energies of HSP90-α ligands for the 2015 D3R grand challenge. Biorg Med Chem. 2016;24(20):4890–9. doi:https://doi.org/10.1016/j.bmc.2016.07.044.

26. Mey ASJS, Jiménez JJ, Michel J. Impact of domain knowledge on blinded predictions of binding energies by alchemical free energy calculations. J Comput Aided Mol Des. 2018;32(1):199–210. doi:10.1007/s10822-017-0083-9. PubMed PMID: PMC5767197.

27. Jiao X, Kopecky DJ, Liu J, Liu J, Jaen JC, Cardozo MG, et al. Synthesis and optimization of substituted furo[2,3-d]-pyrimidin-4-amines and 7H-pyrrolo[2,3-d]pyrimidin-4-amines as ACK1 inhibitors. Bioorg Med Chem Lett. 2012;22(19):6212–7. doi:10.1016/j.bmcl.2012.08.020. PubMed PMID: 22929232; PubMed Central PMCID: PMCUB201.

28. Mahajan K, Mahajan NP. Shepherding AKT and androgen receptor by Ack1 tyrosine kinase. J Cell Physiol. 2010;224(2):327–33. doi:10.1002/jcp.22162.

29. Chua BT, Lim SJ, Tham SC, Poh WJ, Ullrich A. Somatic mutation in the ACK1 ubiquitin association domain enhances oncogenic signaling through EGFR regulation in renal cancer derived cells. Mol Oncol. 2010;4(4):323–34. doi:10.1016/j.molonc.2010.03.001.

30. Molecular Operating Environment (MOE) 2009.1. Chemical Computing Group Inc., 1010 Sherboke St. West, Suite #90, Montreal, QC, Canada, H3A 2R7, 2013. 2013.08 ed.

31. Lougheed JC, Chen RH, Mak P, Stout TJ. Crystal structures of the phosphorylated and unphosphorylated kinase domains of the Cdc42-associated tyrosine kinase ACK1. J Biol Chem. 2004;279(42):44039–45. doi:10.1074/jbc.M406703200. PubMed PMID: 15308621.

32. Case DA, Babin V, Berryman JT, Betz RM, Cai Q, Cerutti DS, et al. AMBER 14. University of California, San Francisco: University of California, San Francisco; 2014 2014.

33. Wang J, Wolf RM, Caldwell JW, Kollman PA, Case DA. Development and testing of a general amber force field. J Comput Chem. 2004;25(9):1157–74. doi:10.1002/jcc.20035.

34. Jakalian A, Bush BL, Jack DB, Bayly CI. Fast, efficient generation of high-quality atomic charges. AM1-BCC model: I. Method. J Comput Chem. 2000;21(2):132–46. doi:10.1002/(SICI)1096-987X(20000130)21:2<132::AID-JCC5>3.0.CO;2-P.

35. Jakalian A, Jack DB, Bayly CI. Fast, efficient generation of high-quality atomic charges. AM1-BCC model: II. Parameterization and validation. J Comput Chem. 2002;23(16):1623–41. doi:10.1002/jcc.10128.

36. Hornak V, Abel R, Okur A, Strockbine B, Roitberg A, Simmerling C. Comparison of multiple Amber force fields and development of improved protein backbone parameters. Proteins: Struct Funct Bioinform. 2006;65(3):712–25. doi:10.1002/prot.21123.

37. Jorgensen WL, Chandrasekhar J, Madura JD, Impey RW, Klein ML. Comparison of simple potential functions for simulating liquid water. J Chem Phys. 1983;79(2):926–35. doi:doi:http://dx.doi.org/10.1063/1.445869.

38. Ȧqvist J. Ion-water interaction potentials derived from free energy perturbation simulations. The Journal of Physical Chemistry. 1990;94(21):8021–4. doi:10.1021/j100384a009.

39. Darden T, York D, Pedersen L. Particle mesh Ewald: An N log(N) method for Ewald sums in large systems. J Chem Phys. 1993;98(12):10089–92.

40. Miller BR, McGee TD, Swails JM, Homeyer N, Gohlke H, Roitberg AE. MMPBSA.py: An Efficient Program for End-State Free Energy Calculations. Journal of Chemical Theory and Computation. 2012;8(9):3314–21. doi:10.1021/ct300418h.

41. Michel J, Essex JW. Prediction of protein-ligand binding affinity by free energy simulations: assumptions, pitfalls and expectations. J Comput Aided Mol Des. 2010;24(8):639–58. doi:10.1007/s10822-010-9363-3. PubMed PMID: 20509041; PubMed Central PMCID: PMCUB186.

42. Loeffler HH, Michel J, Woods C. FESetup: Automating Setup for Alchemical Free Energy Simulations. J Chem Inf Model. 2015;55(12):2485–90. doi:10.1021/acs.jcim.5b00368. PubMed PMID: 26544598; PubMed Central PMCID: PMCUB185.

43. Maier JA, Martinez C, Kasavajhala K, Wickstrom L, Hauser KE, Simmerling C. ff14SB: Improving the Accuracy of Protein Side Chain and Backbone Parameters from ff99SB. Journal of Chemical Theory and Computation. 2015;11(8):3696–713. doi:10.1021/acs.jctc.5b00255.

44. Calabro G, Woods CJ, Powlesland F, Mey AS, Mulholland AJ, Michel J. Elucidation of Nonadditive Effects in Protein-Ligand Binding Energies: Thrombin as a Case Study. J Phys Chem B. 2016;120(24):5340–50. doi:10.1021/acs.jpcb.6b03296. PubMed PMID: 27248478; PubMed Central PMCID: PMCUB191.

45. Chow K-H, Ferguson DM. Isothermal-isobaric molecular dynamics simulations with Monte Carlo volume sampling. Comput Phys Commun. 1995;91(1):283–9. doi:http://dx.doi.org/10.1016/0010-4655(95)00059-O.

46. Åqvist J, Wennerström P, Nervall M, Bjelic S, Brandsdal BO. Molecular dynamics simulations of water and biomolecules with a Monte Carlo constant pressure algorithm. Chem Phys Lett. 2004;384(4):288–94. doi:http://dx.doi.org/10.1016/j.cplett.2003.12.039.

47. Andersen HC. Molecular dynamics simulations at constant pressure and/or temperature. J Chem Phys. 1980;72(4):2384–93. doi:10.1063/1.439486.

48. Barker JA, Watts RO. Monte Carlo studies of the dielectric properties of waterlike models. Mol Phys. 1973;26(3):789–92. doi:10.1080/00268977300102101.

49. Brown SP, Muchmore SW, Hajduk PJ. Healthy skepticism: assessing realistic model performance. Drug Discovery Today. 2009;14(7):420–7. doi:http://dx.doi.org/10.1016/j.drudis.2009.01.012.

50. Shirts MR, Chodera JD. Statistically optimal analysis of samples from multiple equilibrium states. J Chem Phys. 2008;129(12):124105. doi:10.1063/1.2978177.

51. Mey A, Jimenez JJ, Michel J. Impact of domain knowledge on blinded predictions of binding energies by alchemical free energy calculations. J Comput Aided Mol Des. 2018;32(1):199–210. doi:10.1007/s10822-017-0083-9. PubMed PMID: 29134431; PubMed Central PMCID: PMCPMC5767197.

52. Gajiwala KS, Maegley K, Ferre R, He Y-A, Yu X. Ack1: Activation and Regulation by Allostery. PLoS One. 2013;8(1):e53994. doi:10.1371/journal.pone.0053994.

53. Humphrey W, Dalke A, Schulten K. VMD: visual molecular dynamics. J Mol Graph. 1996;14(1):33–8, 27-8. PubMed PMID: 8744570.

54. Origin (OriginLab, Northampton, MA) OriginLab, Northampton, MA.

55. Hunter JD. Matplotlib: A 2D graphics environment. Comput Sci Eng. 2007;9(3):90–5. doi:Doi 10.1109/Mcse.2007.55. PubMed PMID: WOS:000245668100019.

56. https://github.com/mwaskom/seaborn.

57. König G, Hudson PS, Boresch S, Woodcock HL. Multiscale Free Energy Simulations: An Efficient Method for Connecting Classical MD Simulations to QM or QM/MM Free Energies Using Non-Boltzmann Bennett Reweighting Schemes. Journal of Chemical Theory and Computation. 2014:140303161846003. doi:10.1021/ct401118k. PubMed Central PMCID: PMCUB110.

58. Michel J, Tirado-Rives J, Jorgensen WL. Prediction of the water content in protein binding sites. J Phys Chem B. 2009;113(40):13337–46. doi:10.1021/jp9047456. PubMed PMID: 19754086; PubMed Central PMCID: PMCUB195.

59. Cournia Z, Allen B, Sherman W. Relative Binding Free Energy Calculations in Drug Discovery: Recent Advances and Practical Considerations. J Chem Inf Model. 2017;57(12):2911–37. doi:10.1021/acs.jcim.7b00564. PubMed PMID: 29243483; PubMed Central PMCID: PMCUB224.

60. Williams-Noonan BJ, Yuriev E, Chalmers DK. Free Energy Methods in Drug Design: Prospects of “Alchemical Perturbation” in Medicinal Chemistry. J Med Chem. 2018;61(3):638–49. doi:10.1021/acs.jmedchem.7b00681. PubMed PMID: 28745501; PubMed Central PMCID: PMCUB228.

61. Kopecky DJ, Hao X, Chen Y, Fu J, Jiao X, Jaen JC, et al. Identification and optimization of N3,N6-diaryl-1H-pyrazolo[3,4-d]pyrimidine-3,6-diamines as a novel class of ACK1 inhibitors. Biorg Med Chem Lett. 2008;18(24):6352–6. doi:https://doi.org/10.1016/j.bmcl.2008.10.092.

62. Lougheed JC, Chen R-H, Mak P, Stout TJ. Crystal Structures of the Phosphorylated and Unphosphorylated Kinase Domains of the Cdc42-associated Tyrosine Kinase ACK1. J Biol Chem. 2004;279(42):44039–45.

63. Luccarelli J, Michel J, Tirado-Rives J, Jorgensen WL. Effects of Water Placement on Predictions of Binding Affinities for p38alpha MAP Kinase Inhibitors. J Chem Theory Comput. 2010;6(12):3850–6. doi:10.1021/ct100504h. PubMed PMID: 21278915; PubMed Central PMCID: PMCPMC3029023.

64. Bodnarchuk MS, Viner R, Michel J, Essex JW. Strategies to calculate water binding free energies in protein-ligand complexes. J Chem Inf Model. 2014;54(6):1623– 33. doi:10.1021/ci400674k. PubMed PMID: 24684745.

65. Michel J, Henchman RH, Gerogiokas G, Southey MWY, Mazanetz MP, Law RJ. Evaluation of Host–Guest Binding Thermodynamics of Model Cavities with Grid Cell Theory. Journal of Chemical Theory and Computation. 2014;10(9):4055–68. doi:10.1021/ct500368p.

66. Sindhikara Daniel J, Yoshida N, Hirata F. Placevent: An algorithm for prediction of explicit solvent atom distribution—Application to HIV-1 protease and F-ATP synthase. J Comput Chem. 2012;33(18):1536–43. doi:10.1002/jcc.22984.

67. Sridhar A, Ross GA, Biggin PC. Waterdock 2.0: Water placement prediction for Holo-structures with a pymol plugin. PLoS One. 2017;12(2):e0172743. doi:10.1371/journal.pone.0172743. PubMed PMID: 28235019; PubMed Central PMCID: PMCPMC5325533.

68. Hu B, Lill Markus A. WATsite: Hydration site prediction program with PyMOL interface. J Comput Chem. 2014;35(16):1255–60. doi:10.1002/jcc.23616.

69. Ramsey S, Nguyen C, Salomon-Ferrer R, Walker RC, Gilson MK, Kurtzman T. Solvation thermodynamic mapping of molecular surfaces in AmberTools: GIST. J Comput Chem. 2016;37(21):2029–37. doi:10.1002/jcc.24417. PubMed PMID: 27317094; PubMed Central PMCID: PMCPMC5052087.

70. Lazaridis T. Inhomogeneous Fluid Approach to Solvation Thermodynamics. 1. Theory. The Journal of Physical Chemistry B. 1998;102(18):3531–41. doi:10.1021/jp9723574.

71. Lazaridis T. Inhomogeneous Fluid Approach to Solvation Thermodynamics. 2. Applications to Simple Fluids. The Journal of Physical Chemistry B. 1998;102(18):3542– 50. doi:10.1021/jp972358w.

72. Bayden AS, Moustakas DT, Joseph-McCarthy D, Lamb ML. Evaluating Free Energies of Binding and Conservation of Crystallographic Waters Using SZMAP. J Chem Inf Mod. 2015;55(8):1552–65. doi:10.1021/ci500746d.

