## Supplementary material for "Effect of set up protocols on the accuracy of alchemical free energy calculation over a set of ACK1 inhibitors": SI Figures and Table: S1_table.docx

| Protocol | R^2^ | MUE | Slope |
| --- | --- | --- | --- |
| A | 0.37±0.08 | 1.49±0.08 | 0.5 |
| B | 0.49±0.09 | 0.90±0.08 | 0.8 |
| C | 0.30±0.09 | 1.8±0.1 | 0.5 |
| D | 0.69±0.07 | 0.69±0.06 | 1.1 |

^a^ An upper bound of R^2^ = 0.91 ± 0.03 on achievable predictions may be estimated given assumed experimental uncertainties of 0.4 kcal mol^-1^
