## Supplementary figures and images for "Effect of set up protocols on the accuracy of alchemical free energy calculation over a set of ACK1 inhibitors"

### S1_fig.tif

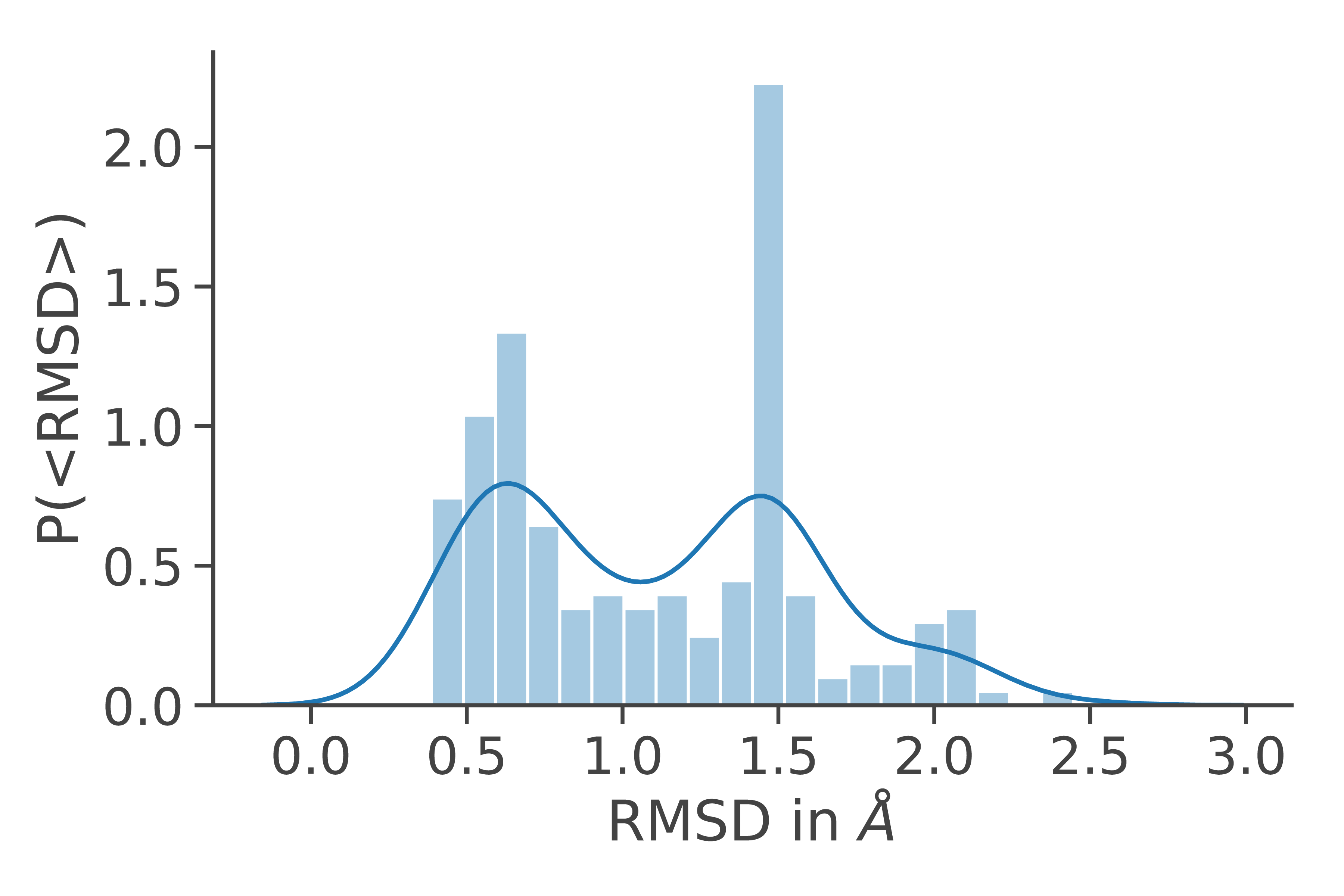

### S2_fig.tif

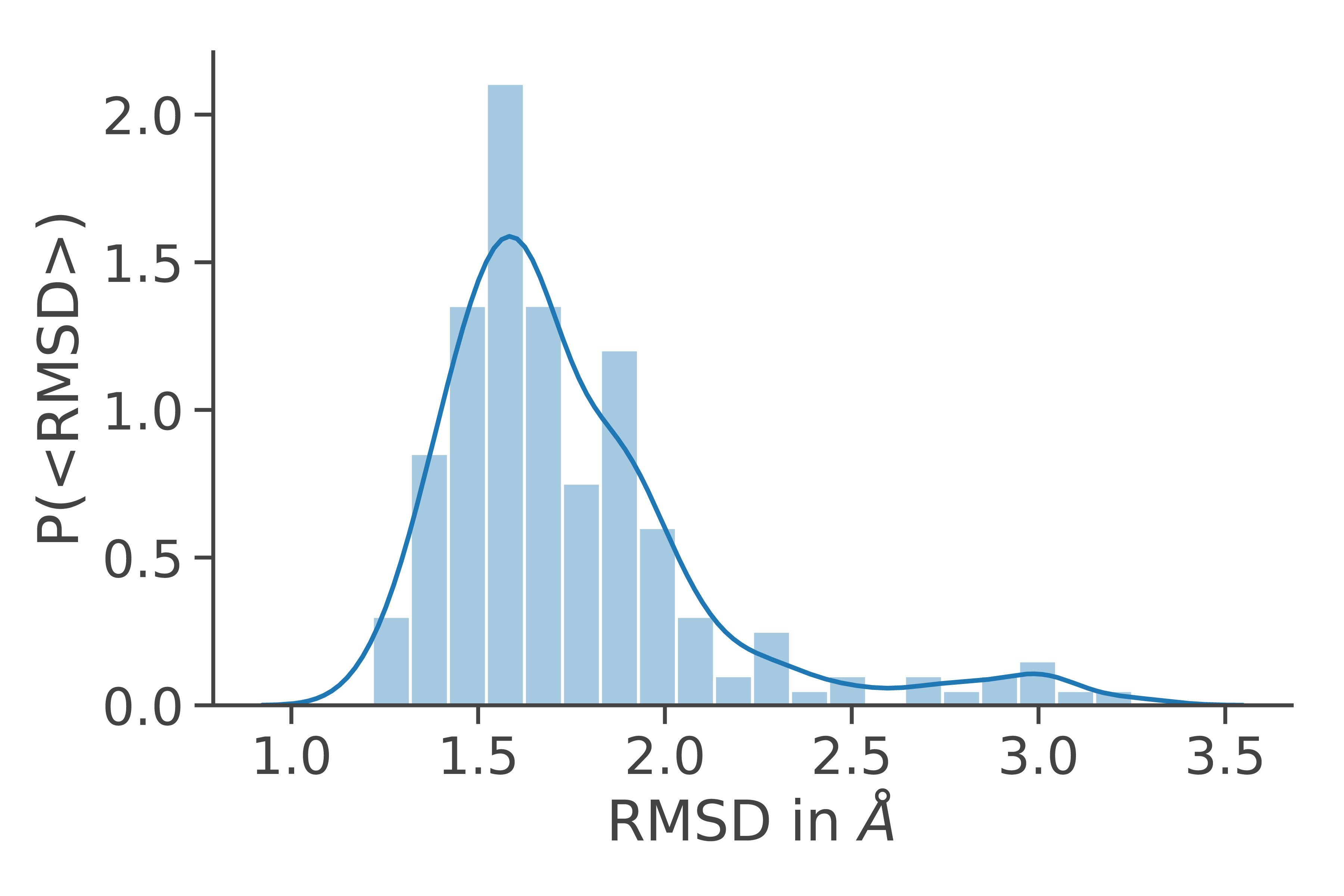

### S3_fig.tif

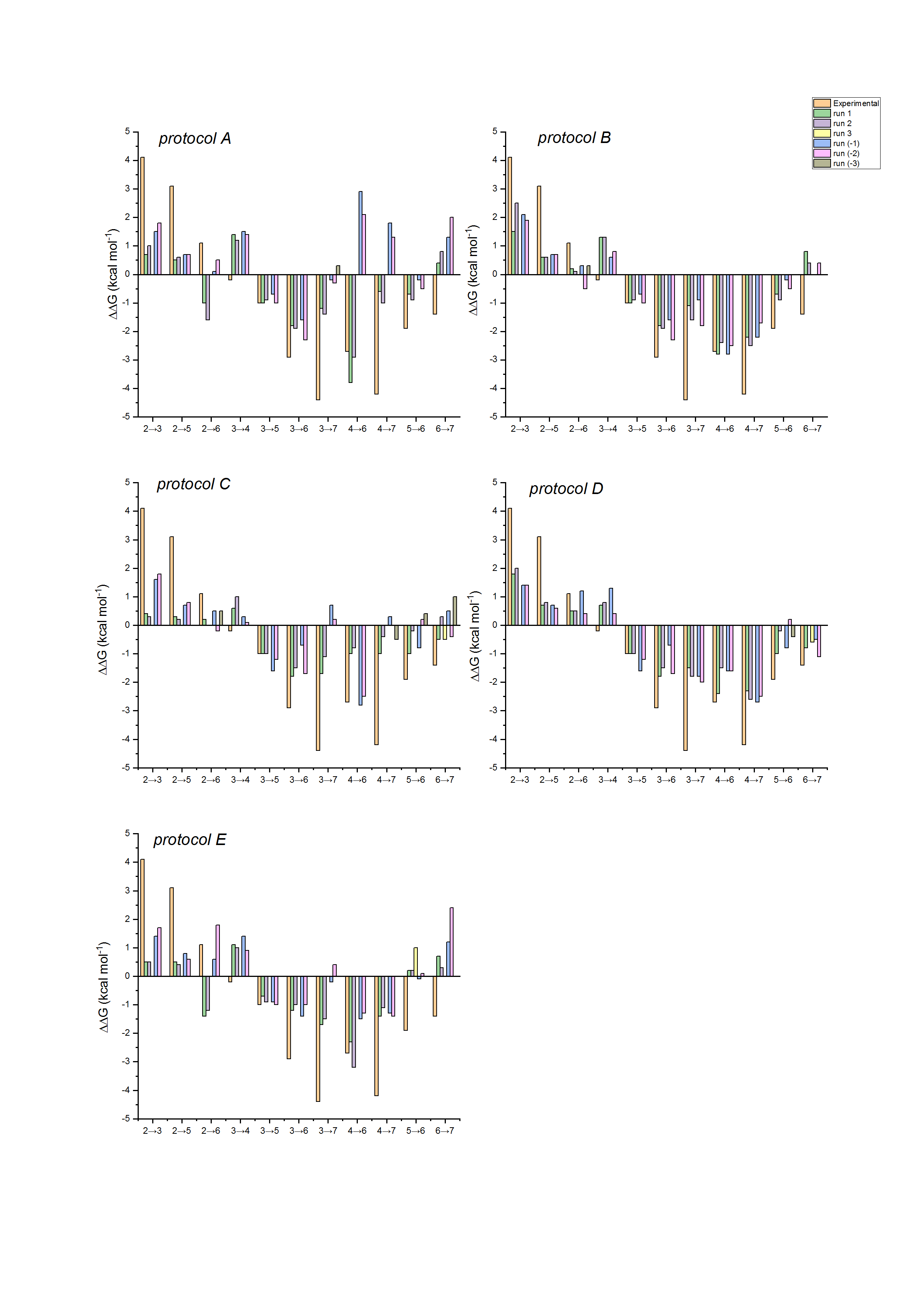

### S4_fig.tif

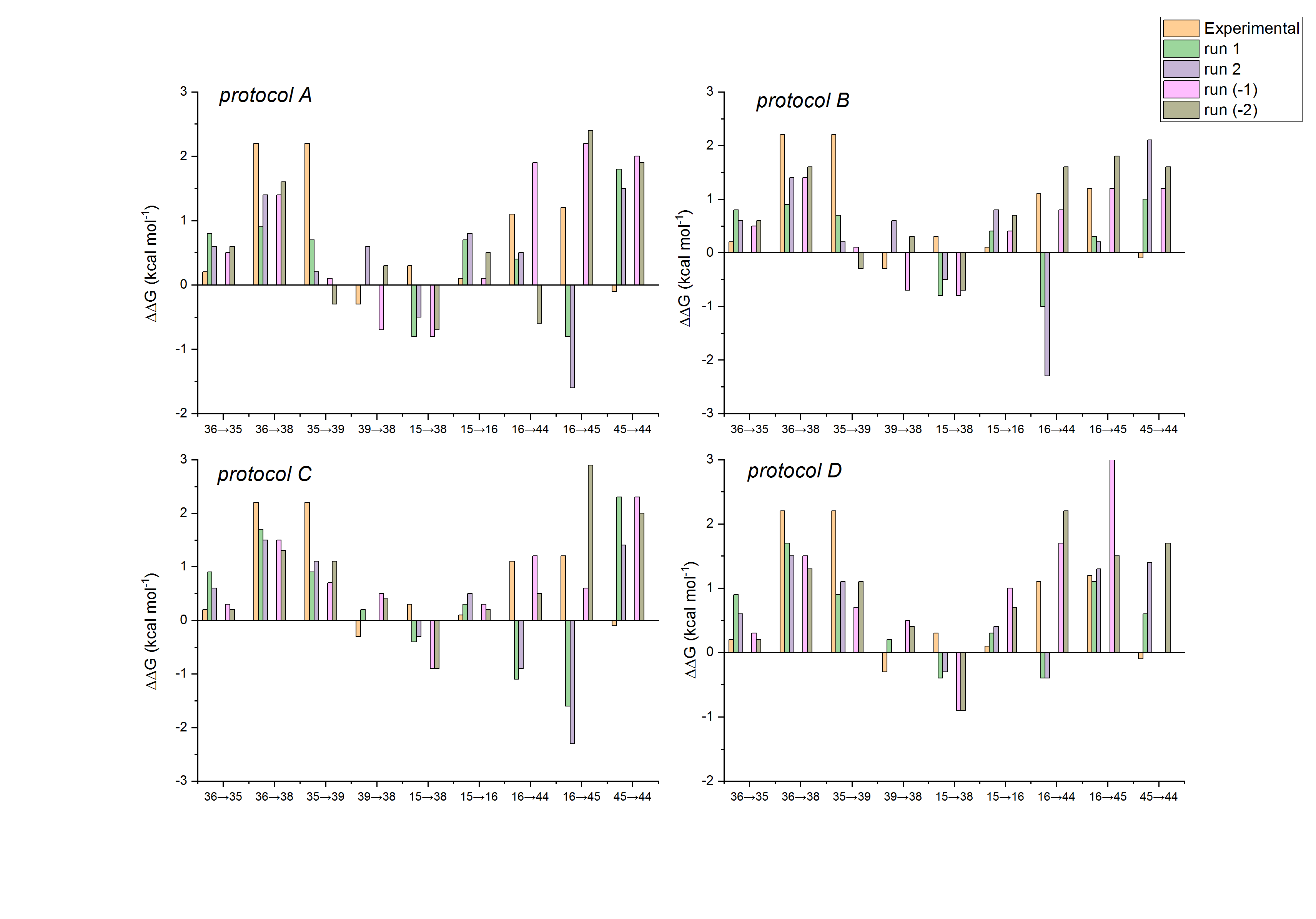

### S5_fig.tif

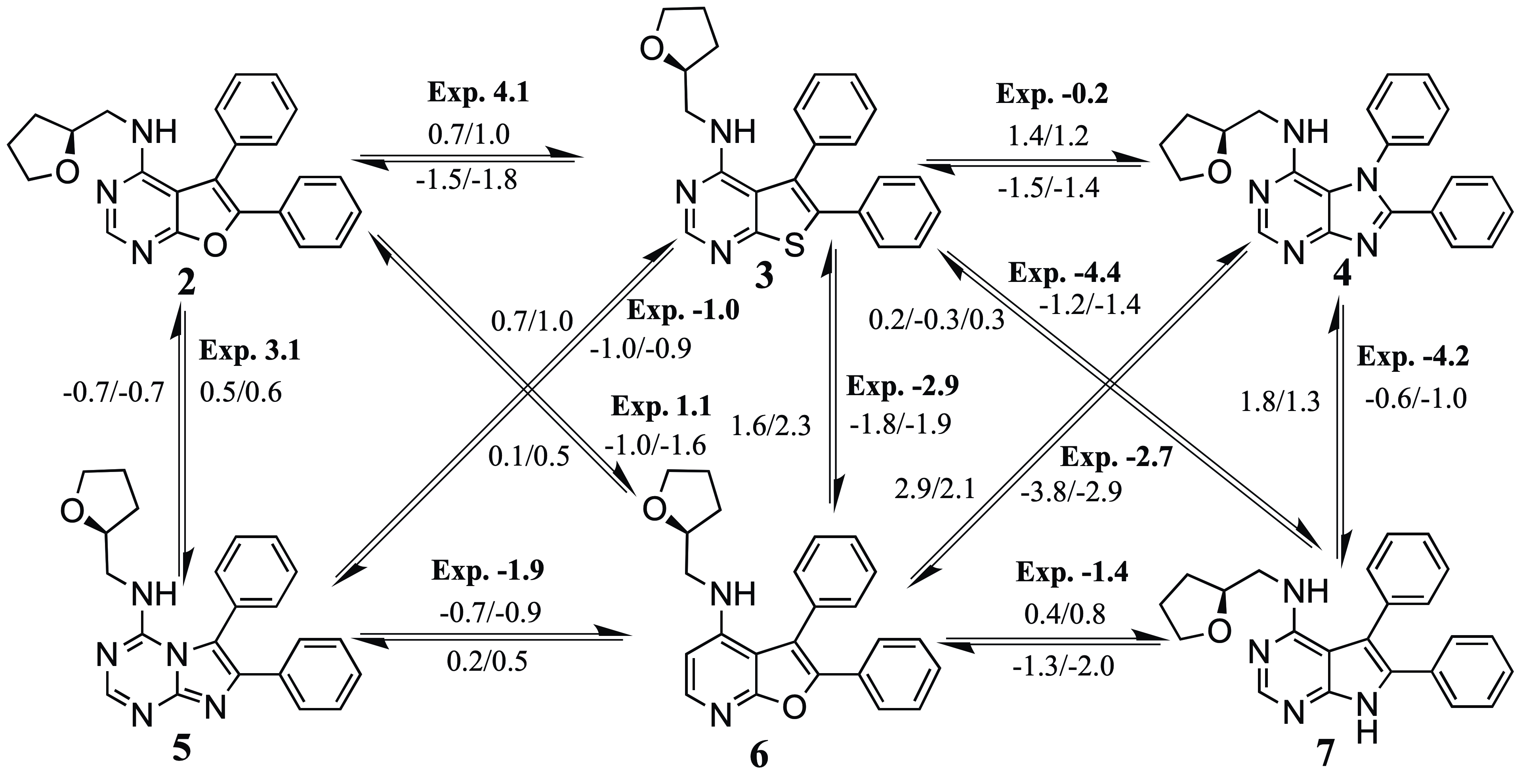

### S6_fig.tif

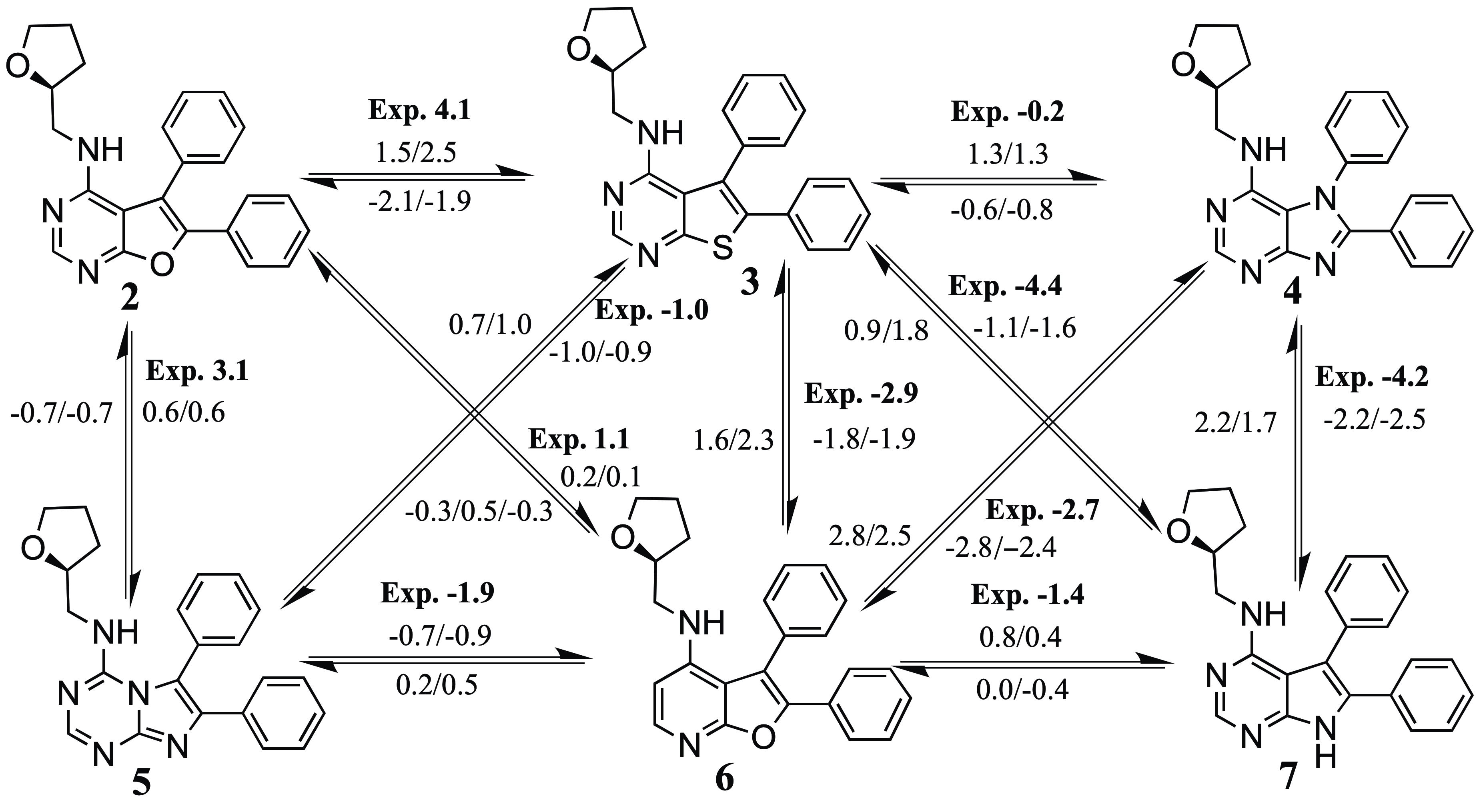

### S7_fig.tif

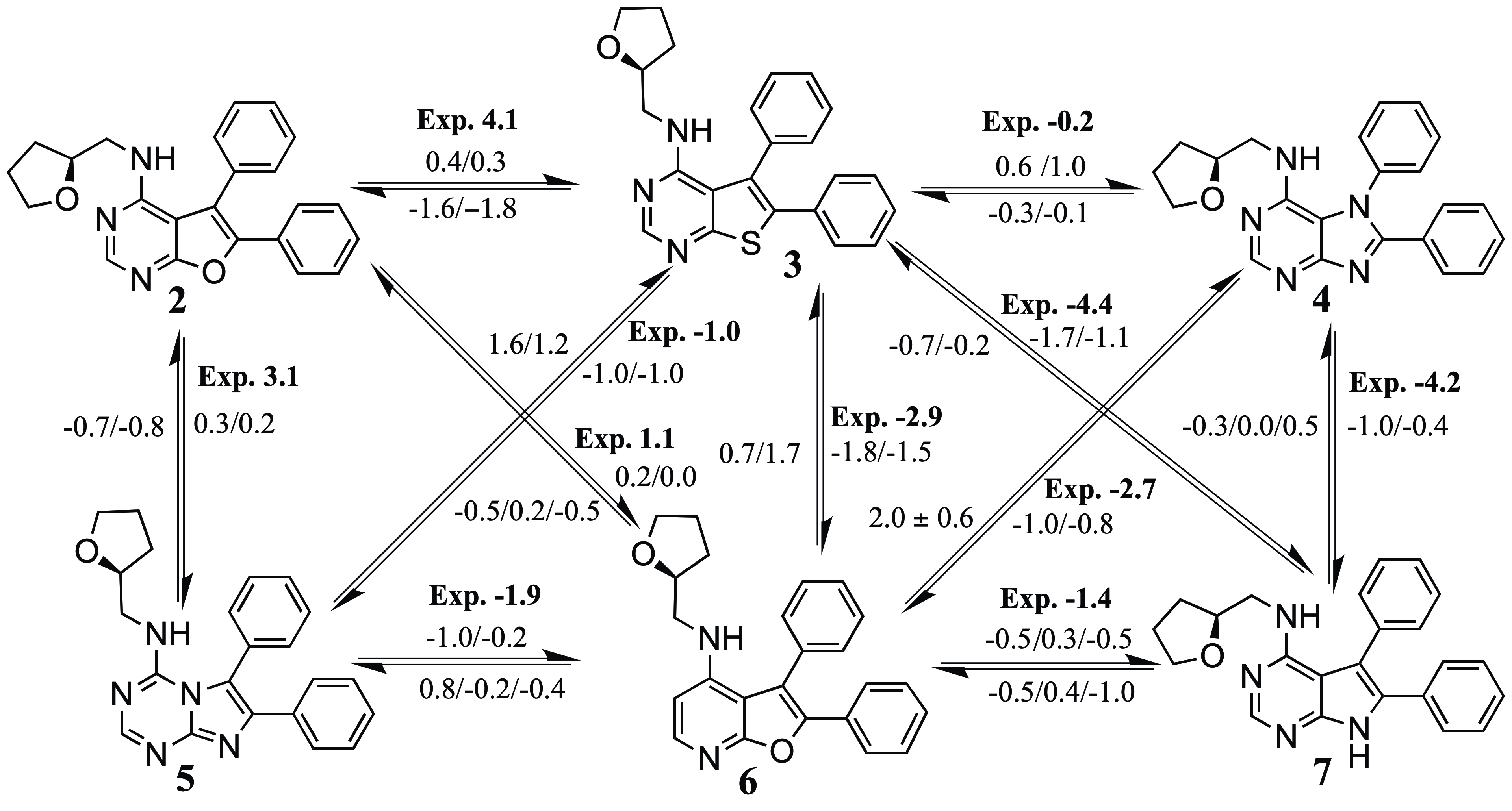

### S8_fig.tif

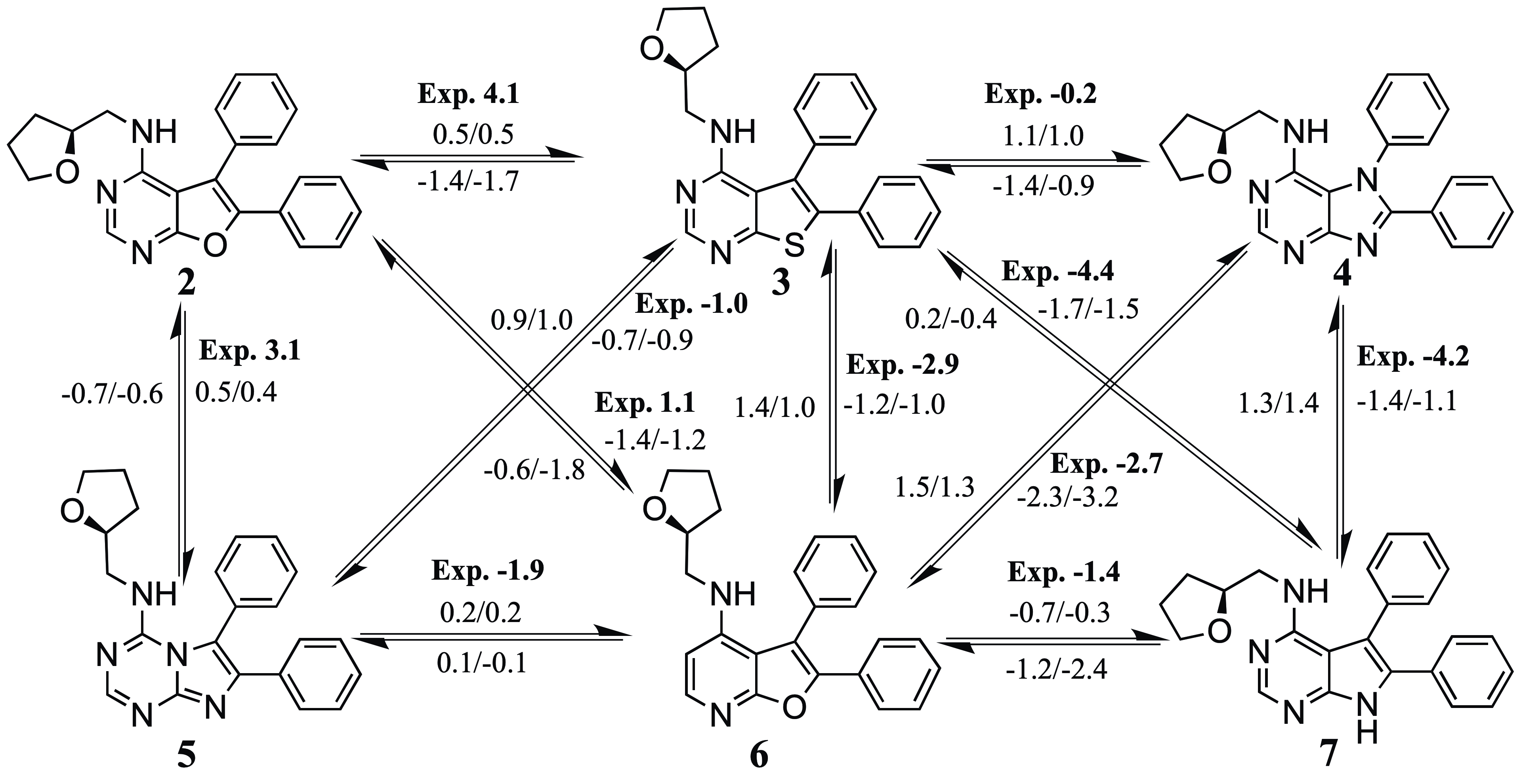

### S9_fig.tif

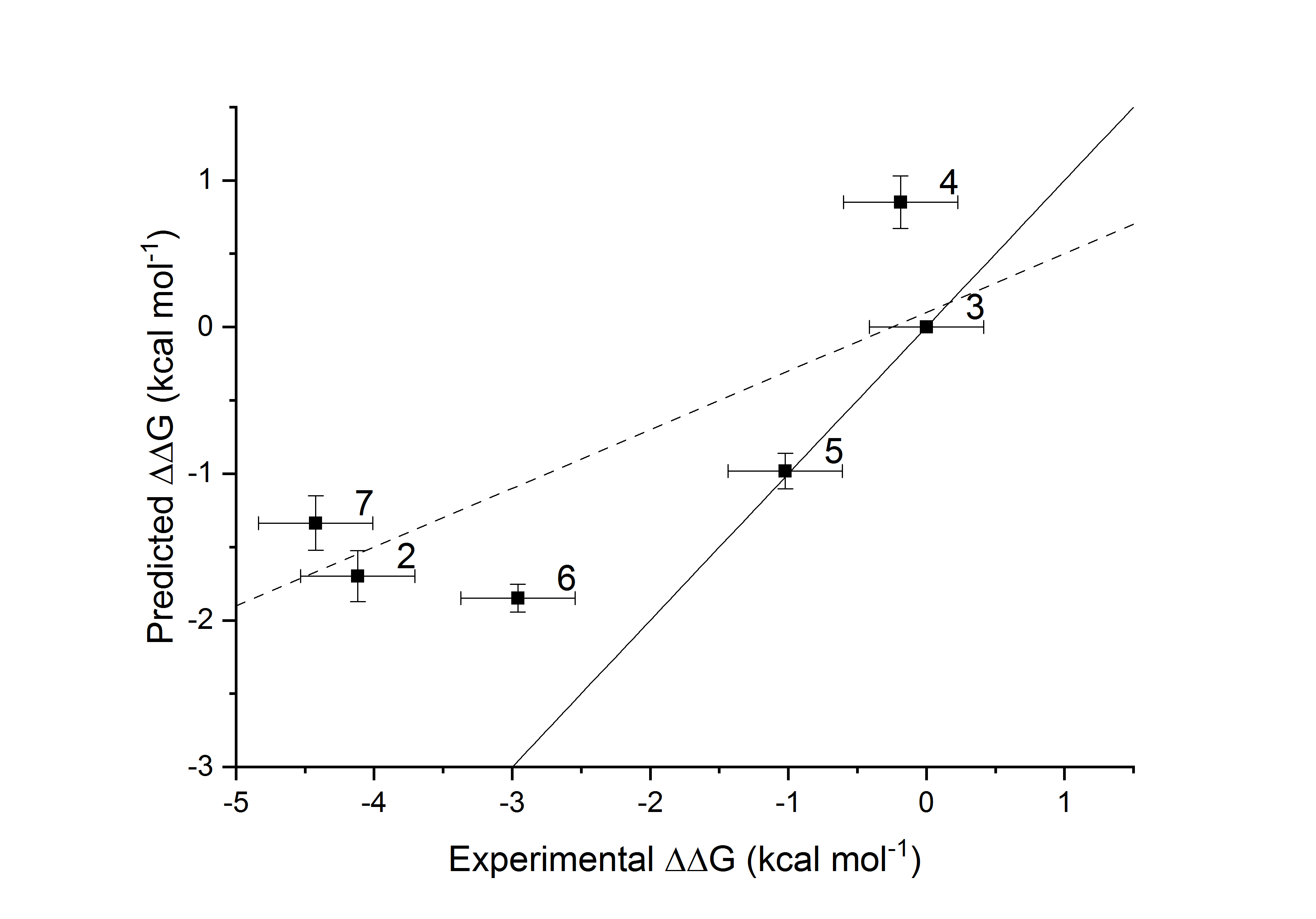

### S10_fig.tif

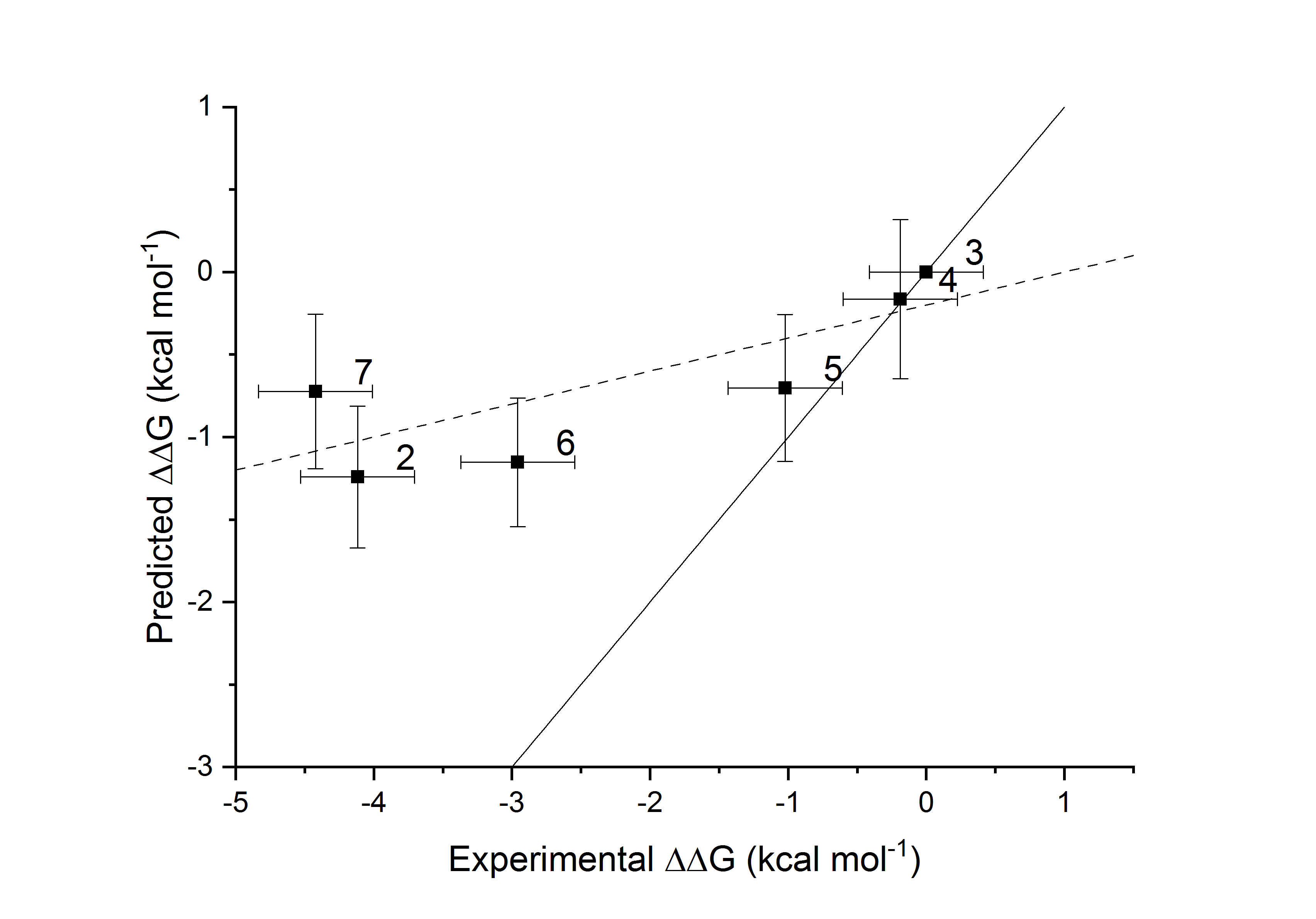

### S11_fig.tif

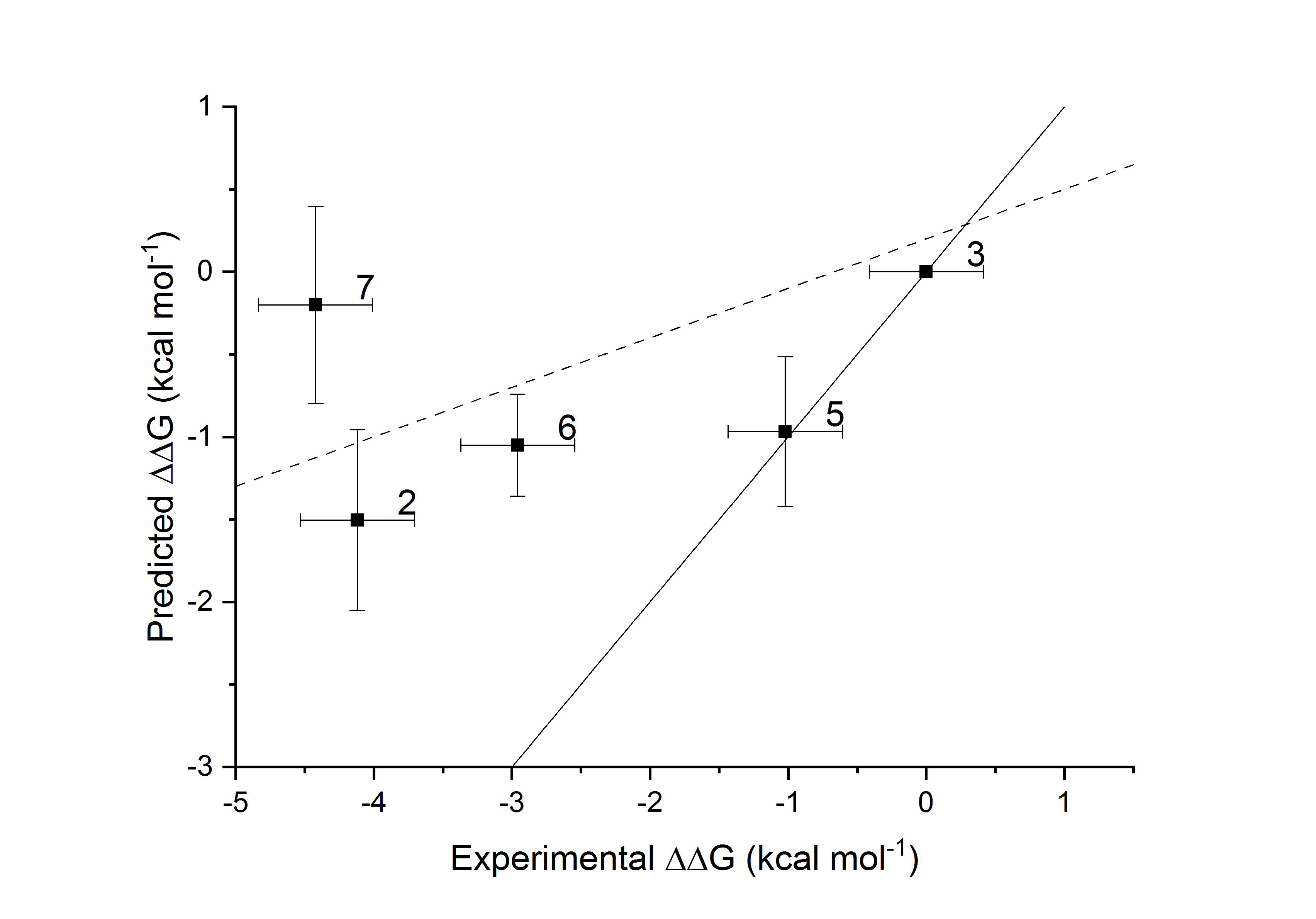

### S12_fig.tif

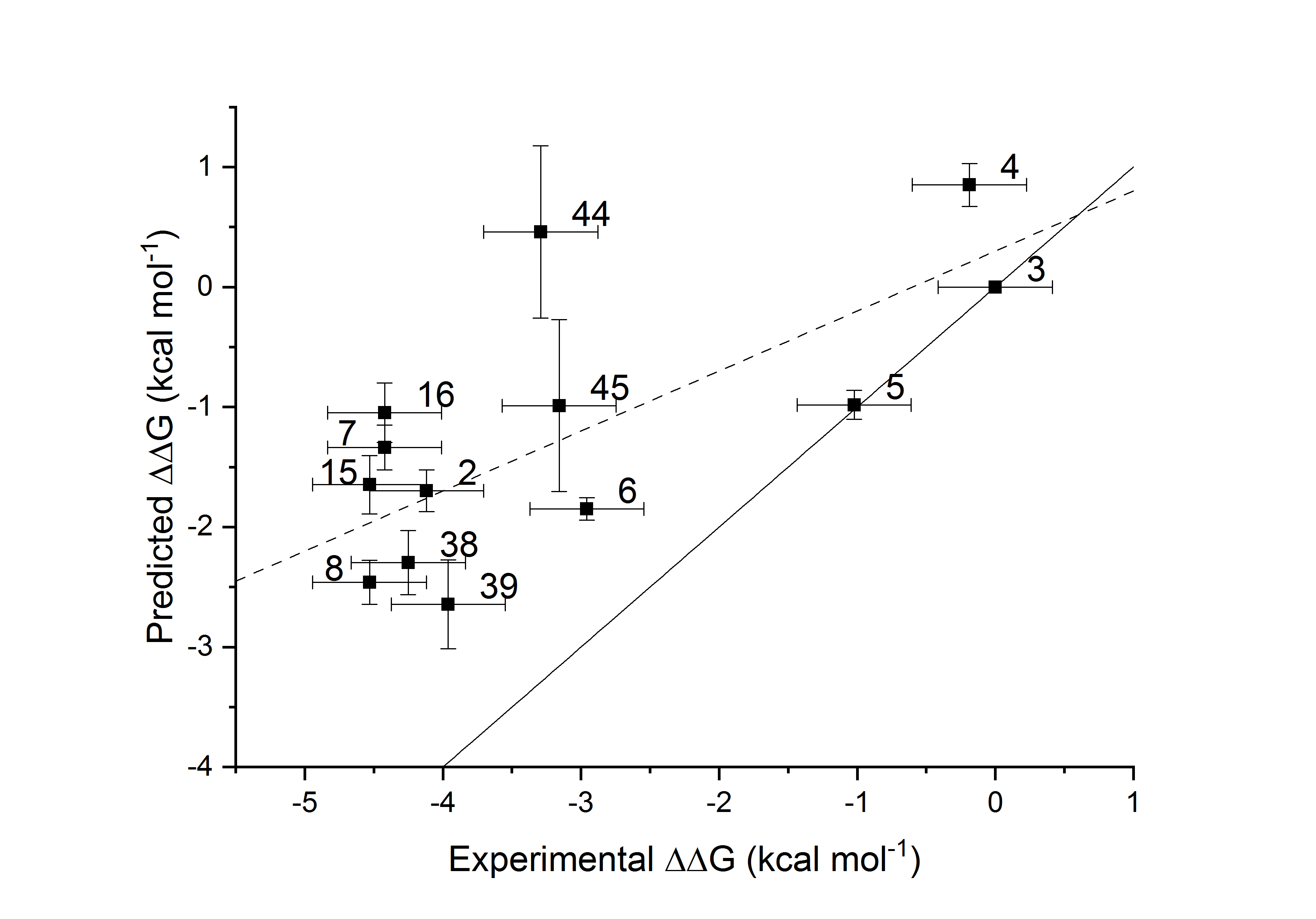

### S13_fig.tif

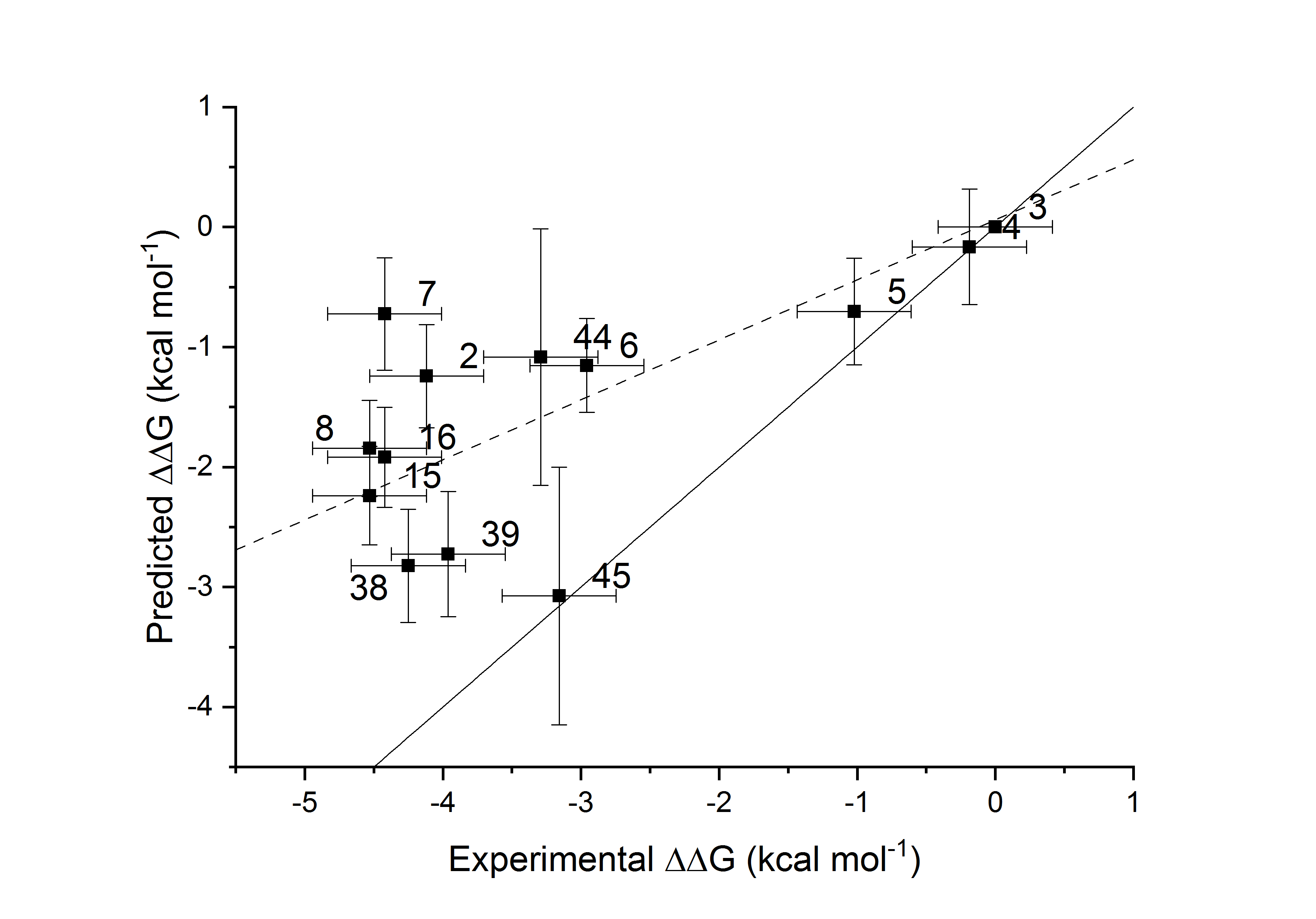

### S14_fig.tif

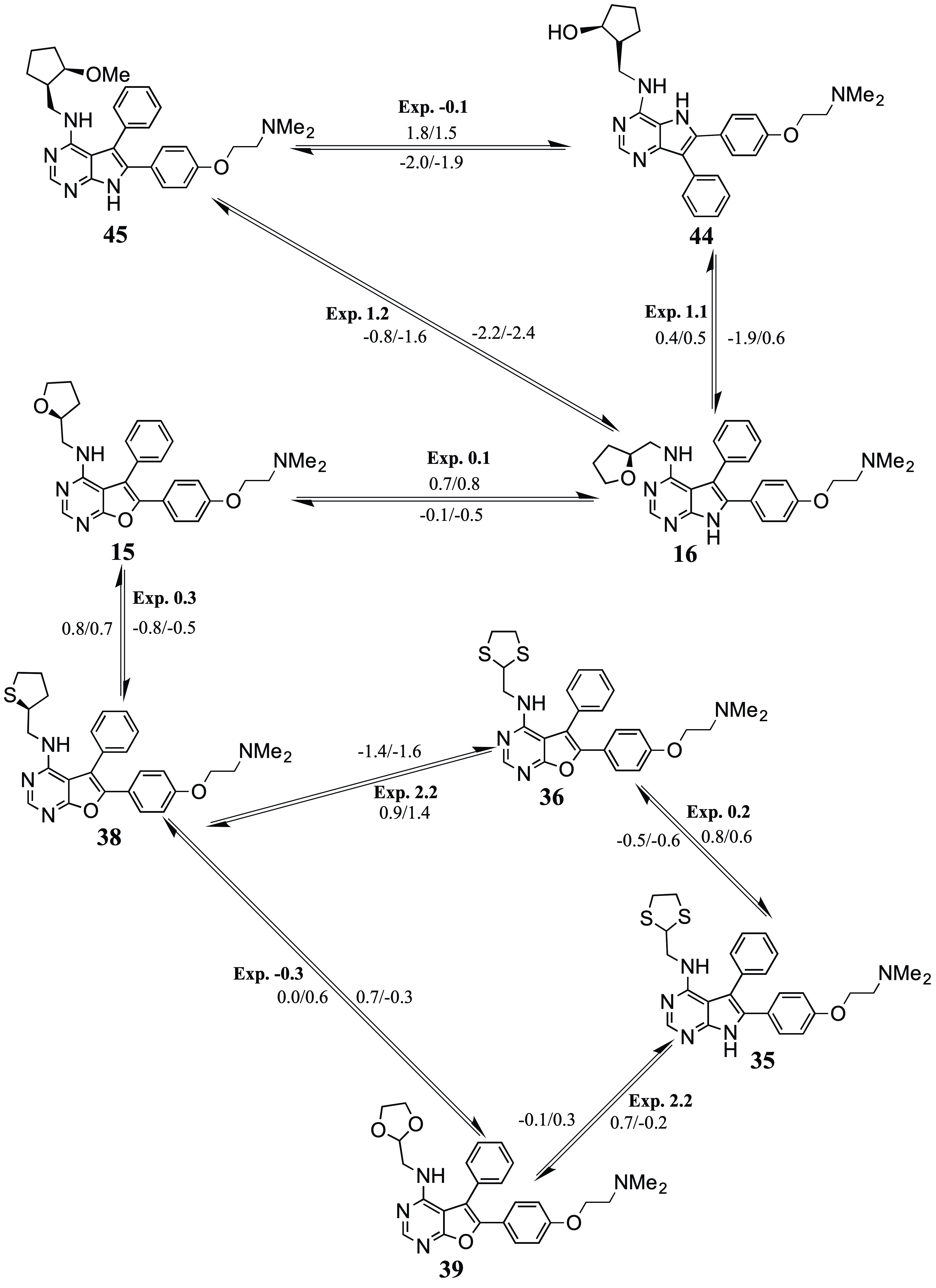

### S15_fig.tif

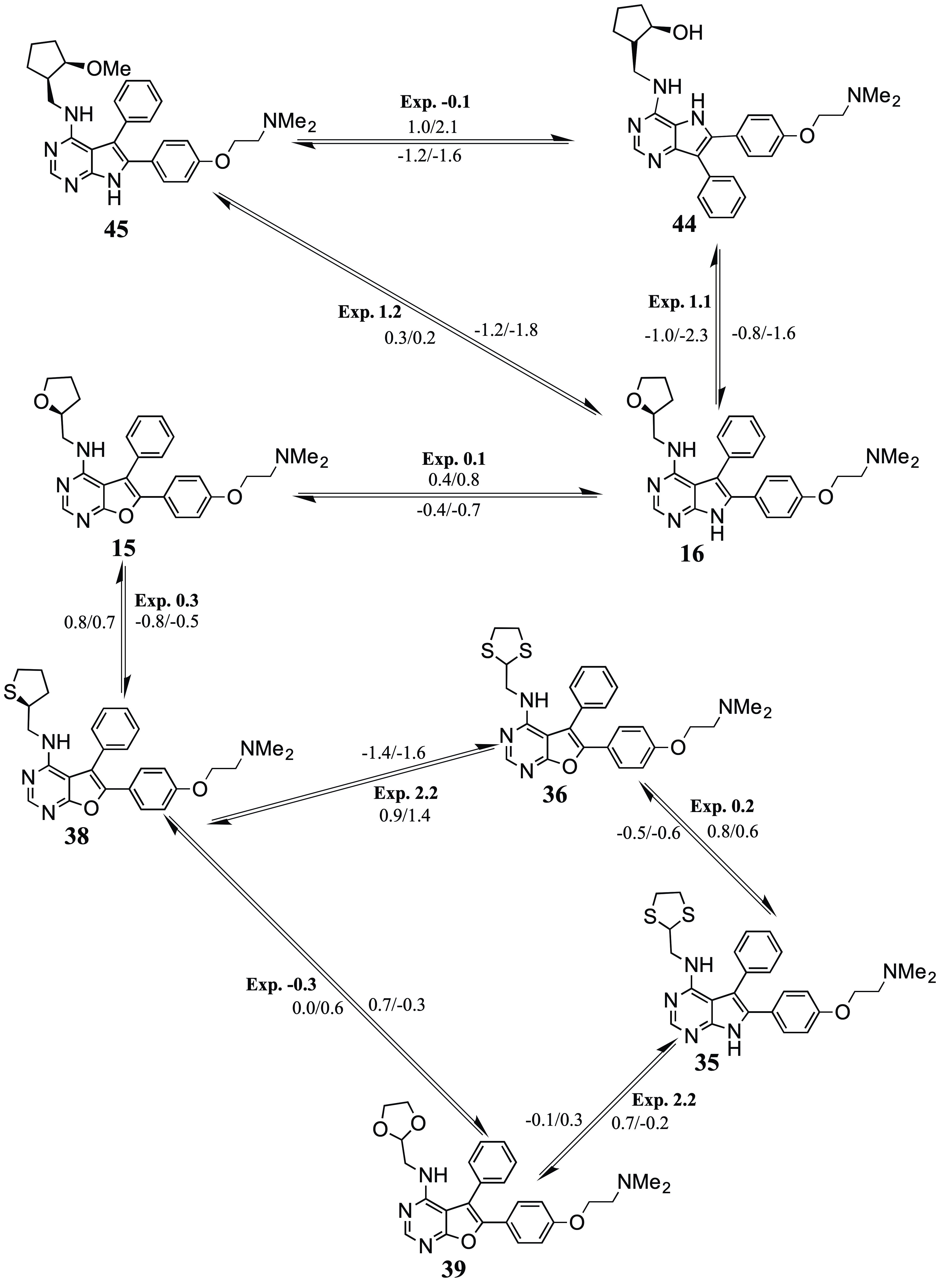

### S16_fig.tif

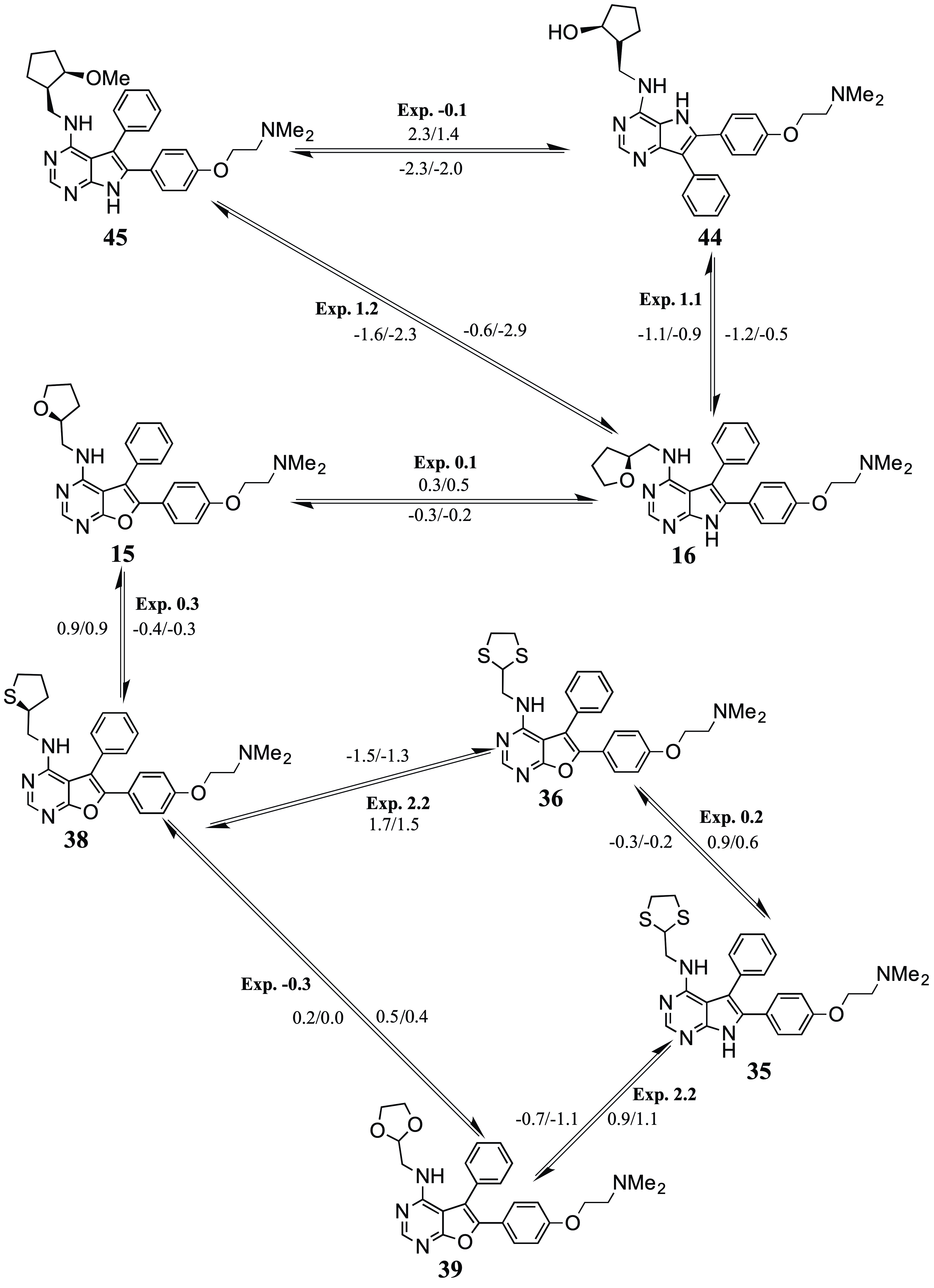

### S17_fig.tif

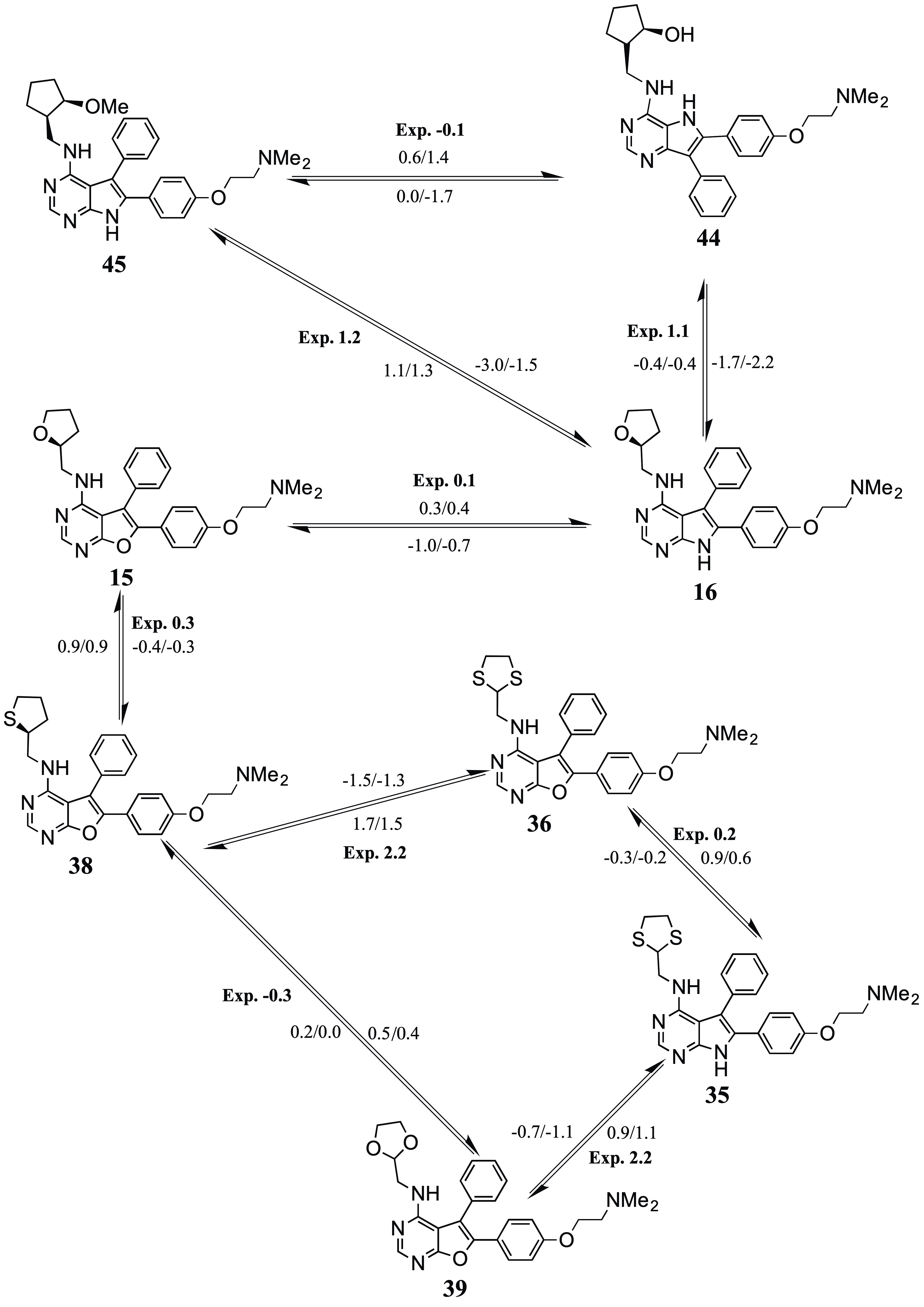

### S18_fig.tif

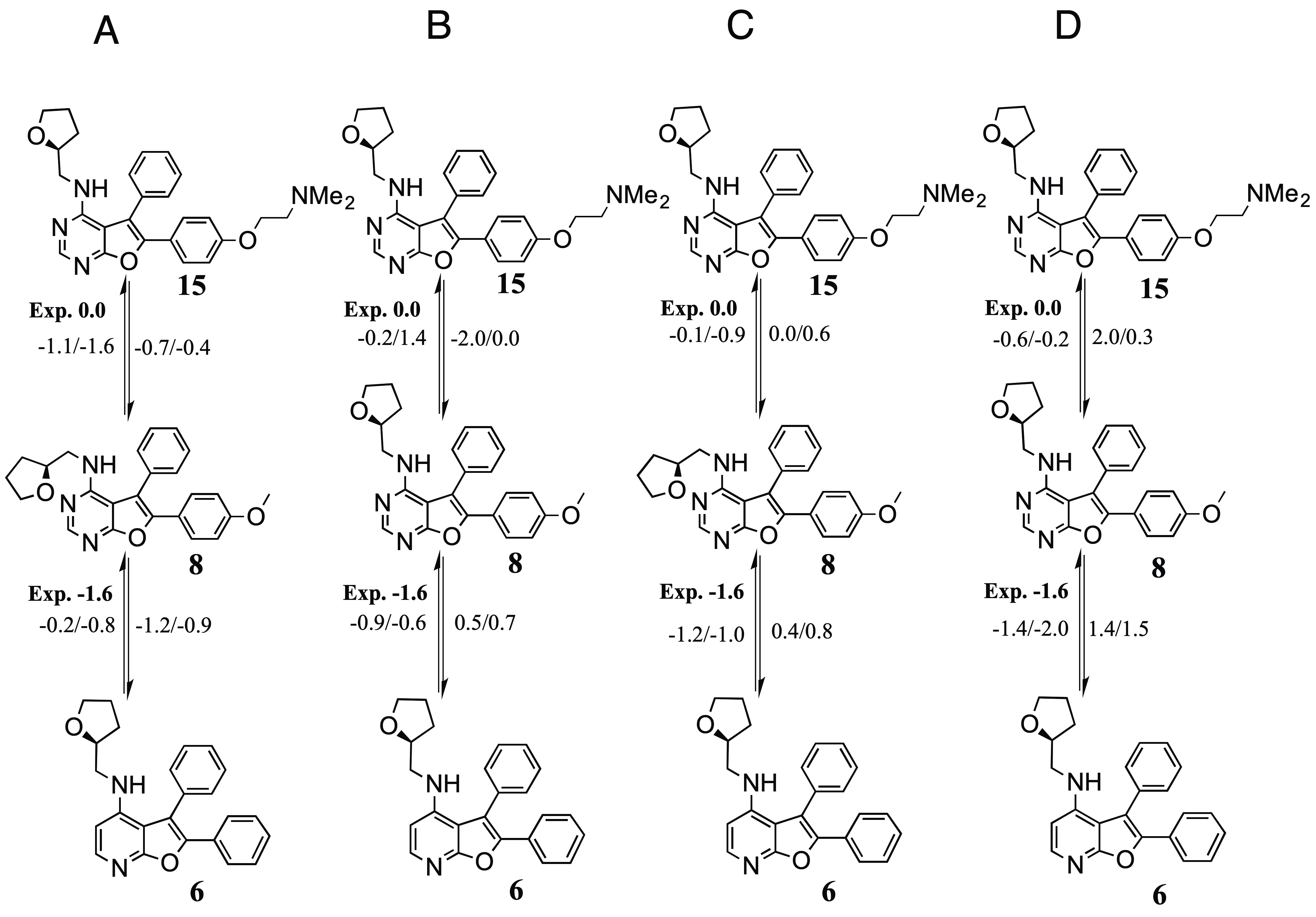
